# A packaging signal-binding protein regulates the assembly checkpoint of integrative filamentous phages

**DOI:** 10.1101/2024.03.19.585677

**Authors:** Ting-Yu Yeh, Michael C. Feehley, Patrick J. Feehley, Vivian Y. Ooi, Yi-Yung Hung, Shao-Cheng Wang, Gregory P. Contreras

## Abstract

Many integrative filamentous phages not only lack Ff coliphage homologues essential for assembly but also have distinct packaging signals (PS). Their encapsidation remains completely uncharacterized to date. Here we report the first evidence of a PS-dependent checkpoint for integrative filamentous phage assembly. Suppressor screening of PS-deficient phages identified an unknown protein, PSB15 (**<u>PS</u>**-**<u>b</u>**inding **<u>15</u>** kDa), crucial for encapsidation. The WAGFXF motif of the PSB15 N-terminus directly binds to PS DNA with conformational change, while suppressor mutations relieve DNA binding specificity constraints to rescue assembly arrest. PSB15 interacts with phospholipid cardiolipin via its basic helix and C-terminus, and recruits PS DNA to the inner membrane (IM). The PSB15-PS complex is released from the IM by interaction between its hydrophobic linker and thioredoxin (Trx), a host protein that is required for Ff assembly but whose mechanisms are still unclear. Live cell imaging shows that thioredoxin and DNA binding regulate the dwelling time of PSB15 at cell poles, suggesting that they both facilitate the dissociation of PSB15 from the IM. Loss of PSB15 or its PS-binding and IM-targeting/dissociation activity compromised virus egress, indicating that the PS/PSB15/Trx complex establishes a regulatory phage assembly checkpoint critical for integrative phage infection and life cycles.

## INTRODUCTION

Assembly and secretion of the archetypical filamentous Ff coliphage was one of the first prokaryotic cell and molecular biology topics studied over 60 years ago (Zinder, 1967; Salivar et al., 1967; Beaudoin and Pratt, 1974; Loh et al., 2019; Mai-Prochnow et al., 2015; Hay and Lithgow, 2019) Virus reproduction requires reasonable efficiency of assembly processes. Virion formation begins with the synthesis of individual coat proteins and is followed by selective binding of the packaging signal (PS) at the inner membrane (IM). PS is a *cis*-acting DNA element that distinguishes viral genomes from cellular DNAs or RNAs (Tate and Peterson, 1974; Shen et al., 1979; Webster et al., 1981; Dotto et al., 1981). During encapsidation, directionality of Ff particles matters as phage particles are structurally asymmetric at their leading and trailing end (Wickner and Killick, 1977; Conners et al., 2023; Jia and Xiang, 2023). At the leading end, pVII and pIX protein complex interact with the PS hairpin of pV-coated phage ssDNA (Russel and Model, 1989; Russel, 1993; Haase et al., 2022). On the phage assembly site, pIV forms a large, gated channel within the outer membrane (OM), while pI and pXI form the IM component of an equimolar, multimeric trans-membrane complex (Ploss and Kuhn, 2010; Gibaud, 2021). As phage DNA traverses through the IM, pV is displaced by the major capsid protein, pVIII, where encapsidation requires machinery made of pI, pXI and pIV (Haase et al., 2022). The order of viral encapsidation with various structural proteins is mostly irreversible with great specificity ( Bruinsma et al., 2021). Nonetheless, across other filamentous phage subtypes, little is known about how the various viral proteins orchestrate a carefully conserved process of infection with host proteins and egress from the membrane.

Integrative filamentous phages play critical roles in human and plant diseases caused by devastating bacteria. These phages integrate their genomes into the host bacterial chromosome directly and replicate like prophages, instead of episomally, like Ff. They have been shown to influence the virulency of their host bacteria, including *Yersinia pestis biovar orientalis* and *Vibrio cholerae*, which respectively causes plague and cholera, and the biofilm formation of *Pseudomonas aeruginosa*, contributing to public health consequences (Waldor and Mekalanos, 1996; Gonzalez et al., 2002; Rice et al., 2009). To date, there are more than 30 different integrative filamentous phages identified from human, animal, and plant pathogens (Mai-Prochnow et al., 2015; Hay and Lithgow, 2019; Yeh et al., 2023). However, very little is known about the PS or encapsidation processes crucial for their replication and infection life cycles.

Thus far, genomic information has shown that integrative filamentous phages of Gram-negative bacteria lack an Ff gIV and gIX homologue. We also verified that Ff gVII homologues are not present in many integrative filamentous phages (Yeh et al., 2023). The absence of these crucial structures prompted us to identify the PS of these phages, what viral (structural and non-structural) factors are involved in their assembly and egress, and how the PS DNA becomes targeted to the IM for encapsidation. To explore the viral assembly of integrative filamentous phages, we first characterized *Xanthomonas* phage PS, which has distinct properties from Ff. Its loop is essential for PS activity, and it contains a shorter (7-8 bp) GC-rich stem compared to f1 (>20 bp). The *Xanthomonas* phage PS also has a different consensus sequence, but importantly, its 5’ to 3’ orientation is not critical for PS competence (Tate and Peterson, 1974; Shen et al., 1979; Webster et al., 1981; Dotto et al., 1981; Russel and Model, 1989; Yeh et al., 2023).

In this study, we found that the PS plays a crucial role in phage assembly distinct from Ff. We identified an unknown protein, PSB15 (**<u>PS</u>**-**<u>b</u>**inding **<u>15</u>** kDa), through genetic suppressor screening, which rescued assembly arrest caused by loss of PS function. PSB15 directly binds to and targets PS and phage DNA to the IM, where this interaction is later released by thioredoxin (Trx), a host protein. We report the first evidence of a filamentous phage assembly checkpoint controlled by PS/PSB15/Trx. These results reveal the underappreciated diversity of the PS in filamentous phages, and demonstrates the many ways filamentous phages can assemble and reproduce, thus redefining the model of integrative filamentous phage assembly.

## RESULTS

### Suppressor screening of PS-interacting proteins

To identify protein factor(s) involved in assembly, we first modified Russel and Model’s method of suppressor screening for PS-interacting protein(s) based on phage plaque morphology (Supplemental Figure S1) (Russel and Model, 1989). We generated a ϕLf-UK phage synonymous T340A/C341G mutant that lost its PS activity but kept gII protein function to facilitate phage DNA replication (Figure 1A and 1C) (Yeh et al., 2023). The ϕLf-UK T340A/C341G plaque morphology was tiny (0.1-0.2 mm) compared to the wild-type phage (1-1.2 mm). On the bacterial lawn, normal-sized plaques could be found with a high frequency of 5×10^-2^ to 10^-4^. The normal-sized plaques were collected to amplify phages. These phage mutants represented either true reversion (A340 → T340) or second-site mutations (pseudorevertants), which contain suppressor mutations able to compensate for the loss of PS activity.

**Figure 1.**
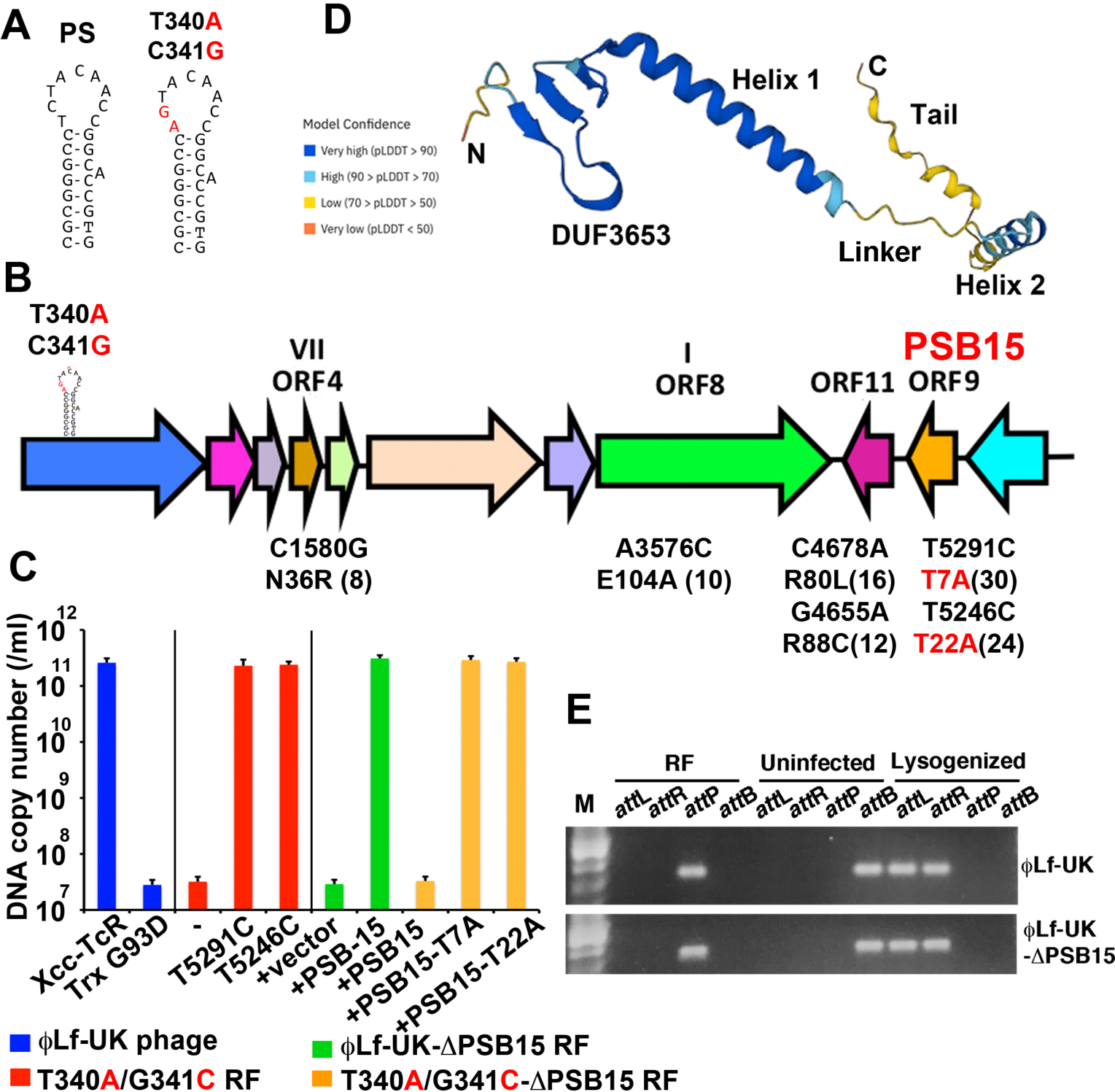
Identification of PSB15, a phage-encoded protein factor essential for assembly. (A) ϕLf-UK PS T340A/C341G mutant was used for suppressor screening, whose procedures are described in detail in Supplemental Figure S1. (B) Summary of suppressor mutations. Numbers of each isolated suppressor are shown in parentheses (total 100 sequences). Suppressor T5291C (Thr7Ala) and T5246C (Thr22Ala) mutations in PSB15 are labeled in red. (C) (Blue) Phage yields of ϕLf-UK infection in Xcc-TcR and Xcc-TcR-TrxA G93D bacteria. (Red) ϕLf-UK-T340A/C341G and its suppressor T5291C and T5246C mutants released in the medium were measured using qPCR. (Green and organe) *In vivo* complement assay was carried out by pPSB15 plasmid, which encodes PSB15 protein. pPSB15 was co-transformed with ϕLf-UK-ΔPSB15 (green) and T340A/C341G-ΔPSB15 RF (orange), followed by qPCR to measure phage production. (D) AlphaFold prediction of PSB15 structure (UniProt: Q8P905). Model confidence based on a per-residue confidence score (pLDDT) is shown by color. (E) RF replication (*att*P) and site-specific integration of ϕLf-UK-ΔPSB15 phage DNA (*att*L and *att*R) into the host chromosome was analyzed by PCR. The PCR primers, detail methods, and ϕLf-UK integration (upper) were described previously (24).

We added a second screening step to find suppressors by using these phage mutants as helper viruses to infect bacteria transformed with pORIPS-PS(-) plasmid. When infected with ϕLf-UK helper phage, pORIPS-PS(-) contains an antisense PS [PS(-)] sequence unable to assemble transducing particles (TPs) efficiently (0.9-1.5% yield compared to the canonical PS) (24). Pseudorevertants, but not true revertants, can provide suppressors *in trans* to compensate for the deficient production of pORIPS-PS(-) TP. Therefore, pseudorevertants can be screened by detecting the significant increase of pORIPS-PS(-) TPs released into the medium using qPCR. A total of 123 pseudorevertants were isolated from 4 independent screens (1026 phages with normal-sized plaques). The mutations of 100 suppressors were mapped by whole genome sequencing, and the results are summarized in Figure 1B.

Our data showed that gVII, gI, and two unknown genes, ORF9 and ORF11, genetically interact with the PS. These pseudorevertants all contain a single nonsynonymous mutation. A total of 30 and 24 pseudorevertants contained the substitution of T5291C and T5246C, respectively, which encode a Thr7Ala (T7A) and Thr22Ala (T22A) mutation in the ϕLf-UK ORF9 protein (here named as **<u>PS</u> <u>b</u>**inding **<u>15</u>** kDa, PSB15). C4678A and G4655A mutations in ORF11 (Arg80Leu and Arg88Cys in protein) were identified among 16 and 12 pseudorevertants, respectively. We also isolated 8 pseudorevertants with C1580G (ORF4/gVII Asn36Arg) and 10 ones with A3576C (ORF8/gI Glu104Ala) mutations. Because ORF9/PSB15 is an unknown protein and conserved among all *Xanthomonas* integrative filamentous phages, we chose to focus on the encapsidation roles of ORF9/PSB15 in this study.

### PSB15 protein is required for viral assembly *in trans* but not phage DNA integration or replication

T340A/C341G mutations cause a drastic decrease in phage reproduction yield (∼10^-4^ fold), measured by phage particle DNA copy numbers released in the medium (Figure 1C, red). Suppressors T5291C (PSB15-T7A) and T5246C (PSB15-T22A) can rescue this assembly arrest phenotype, consistent with TP production results in our screening. PSB15 is conserved among *Xanthomonas* and *Stenotrophomonas* filamentous phages (24) (Figure 2D). However, its function has never been reported. Based on AlphaFold algorithm predictions, PSB15 contains 5 putative structure domains: N-terminal DUF3653 (AA 1-29), helix 1 (AA 30-58), linker (AA 59-68), helix 2 (AA 69-81), and a disordered C-terminal tail (AA 82-108) (Figure 1D).

**Figure 2.**
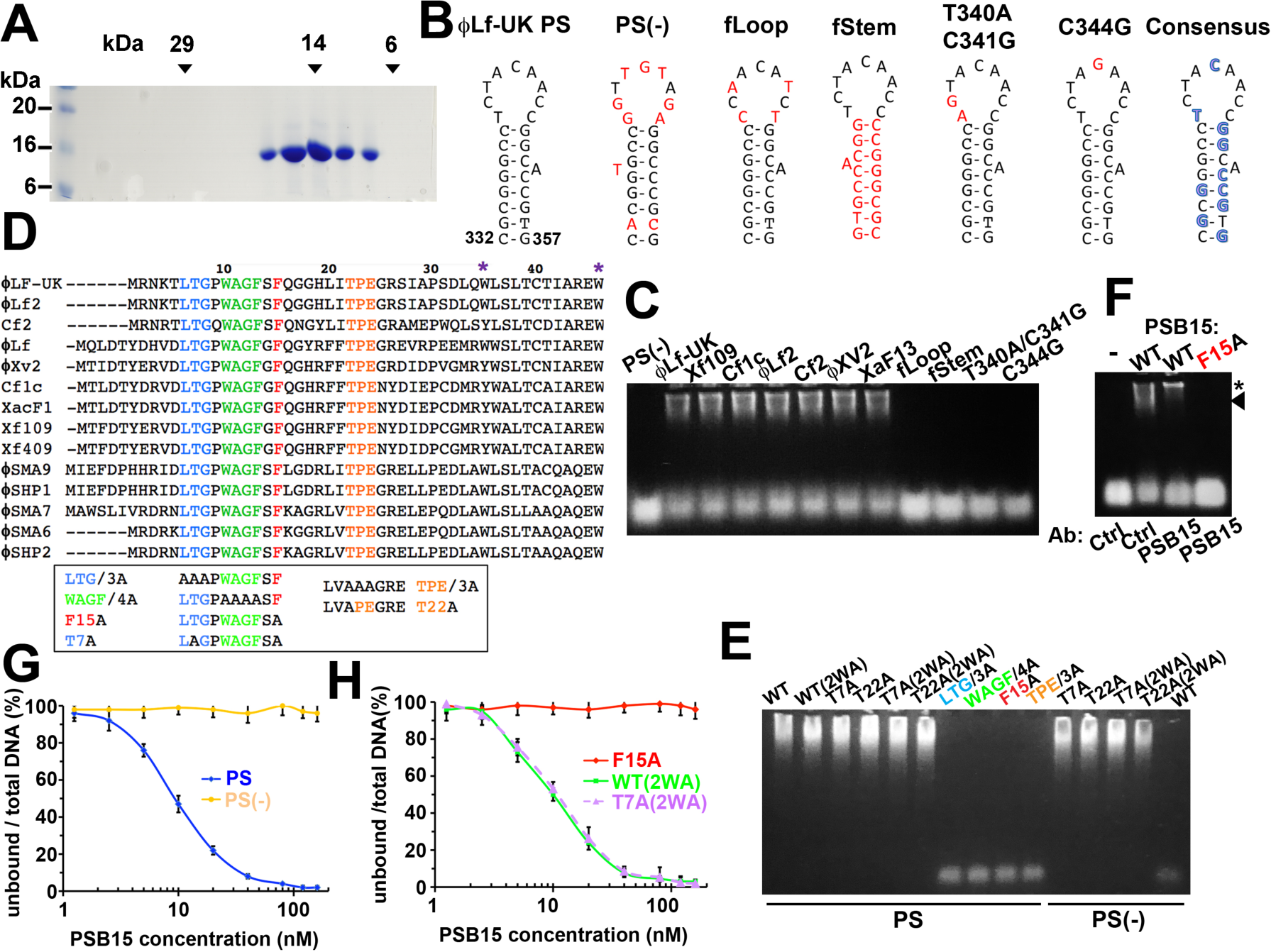
DNA binding properties of PSB15. (A) Gel filtration chromatography of recombinant PSB15. The peak fractions were collected and analyzed by 13.5% SDS-PAGE and Coomassie blue staining. (B and C) Oligonucleotides of ϕLf-UK PS and its mutant (B), and *Xanthomonas* filamentous phages PS (24) were used for EMSA (C). PS DNA-PSB15 complex was separated in 1.5% agarose gel and stained for SYBR Gold. The free DNA probe is at the bottom. (D) Sequence alignment of PSB15 N-terminus of *Xanthomonas* and *Stenotrophomonas* filamentous phages. Mutations are listed in the box. (E) EMSA of PSB15 mutants with ϕLf-UK PS and PS (-). PSB15 tryptophan 35 and 46 [purple asterisks in (D)] were replaced with alanine (2WA). The free DNA probe is at the bottom. (F) PS-PSB15 complex (arrowhead, F) and a supershift by PSB15 antibody (asterisk, F) were shown. (G and H) PSB15-PS DNA binding kinetics in (C) and (E) from 3 independent experiments.

To confirm that the ORF9 product is a protein, rather than a non-coding RNA, and that it engages in encapsidation, the start codon of ORF9 was mutated in replicative form (RF) DNA (ϕLf-UK-ϕPSB15) and transformed into bacteria. The PSB15 protein expression in ϕLf-UK-ϕPSB15-transformed bacteria can not be detected (Figure S2). ϕLf-UK-ϕPSB15 phage reproduction was significantly reduced, and the plaque size was tiny (Figure 1C), indicating that loss of PSB15 phenocopies ϕLf-UK PS mutations (Yeh et al., 2024). An *in vivo* complement assay showed that phage production of ϕLf-UK-ϕPSB15 can be fully rescued by co-transformation with pPSB15 plasmid (Figure 1C, green), which ectopically expresses the PSB15 protein (Figure S2). Furthermore, the phage assembly defect caused by the PS mutant, T340A/C341G (T340A/C341G-ϕPSB15 RF transformants), can be rescued by ectopic expression of PSB15-T7A or -T22A, but not wild type (WT) PSB15 (Figure 1C, orange). Taken together, this and later data confirm that PSB15 protein is crucially involved in encapsidation as a *trans*-acting factor, and its amino acid T7A and T22A mutants are suppressors for the PS mutant.

We also found that integration of ϕLf-UK-ϕPSB15 into the host chromosome (*att*L and *att*R), intracellular RF dsDNA replication (*att*P), phage ssDNA, and coat protein synthesis (data not shown) remained unchanged (Figure 1E), indicating that PSB15 is not essential in these functions. Because the PSB15 protein can be detected in ϕLf-UK-infected cell lysate, but not in CsCl-purified virions, it appears not to be a structural or minor coat protein (Figure S2).

### PSB15 is a PS DNA-binding protein

Our results suggest interaction between PSB15 and the PS. To test whether PSB15 binds to the PS directly, we expressed and purified recombinant PSB15 protein from *Escherichia coli*. Recombinant protein was recovered from the inclusion body with 6M urea and refolded in CAPS buffer with reducing urea concentrations. A single peak of PSB15 protein was obtained in gel filtration chromatography at approximately 15 kDa (Fig. 2A), indicating that soluble PSB15 is a monomer.

We sought to examine PSB15 and PS binding using electrophoretic mobility shift assays (EMSA). Figure 2C shows that PSB15 can form a complex with PS oligonucleotides of all *Xanthomonas* filamentous phages (ϕLf-UK/ ϕLf, Xf109, Cf1c/XacF1, ϕLf2, Cf2, ϕXv2, and XaF13) with similar equilibrium dissociation constant (K_d,app_ =10.2 ± 1.1 to 13.4 ± 2.1 nM, Table S1). PSB15 was unable to bind to PS(-) oligonucleotide (Figure 2C and 2G), poly(A)_25_, poly(T)_25_, poly(C)_25_, or poly(G)_25_ (Table S1), indicating that binding is sequence specific. The specificity of PS-PSB15 complex was also demonstrated by a supershift generated by anti-PSB15 antibody, but not by the control antibody (Figure 2F). No intermediate band suggested a highly cooperative mode of interaction between PSB15 and PS DNA.

Furthermore, PS stem (fStem) and loop mutants (fLoop, T340A/C341G, C344G) failed to bind to PSB15 (Figure 2B and 2C). We also tested other PS mutants listed in Table S1. Our data demonstrate that the consensus sequence for encapsidation [GGX(A/-)CCG(C/T)G in the stem and T340, C344 in the loop] is also essential for the PS binding to PSB15 (Figure 2B).

### DNA binding properties and essential motifs of PSB15

Alignment of the N-terminal DUF3653 domain of *Xanthomonas* and *Stenotrophomonas* filamentous phages reveals three conserved motifs (^6^LTG, ^10^WAGFXF, and ^22^TPE) at a ϕ hairpin structure seen in AlphaFold. Suppressors T7A and T22A are located at the LTG and TPE motif, respectively (Figure 2D and 3A). This formulated the hypothesis that LTG/WAGFXF/TPE is the PS DNA binding motif. To test this, we purified recombinant PSB15 mutants with alanine substitutions (LTG/3A, WAGF/4A, TPE/3A, F15A, T7A, T22A). For comparison, we also prepared PSB15 mutants in other domains (Supplemental Figure S3). EMSA showed that mutations in LTG/3A, WAGF/4A, TPE/3A, and F15A, but not other domains, completely abolished PS DNA binding (Figure 2E, and 5C), suggesting that LTG/WAGFXF/TPE contains the DNA binding motif. Suppressors T7A and T22A showed distinct properties because they were able to bind to PS(-) and canonical PS DNA with similar affinity (Figure 2E and Table S2). These results not only indicate that T7 and T22 contribute to PS DNA binding specificity, but also explain why T7A and T22A can rescue phage assembly arrest.

**Figure 3.**
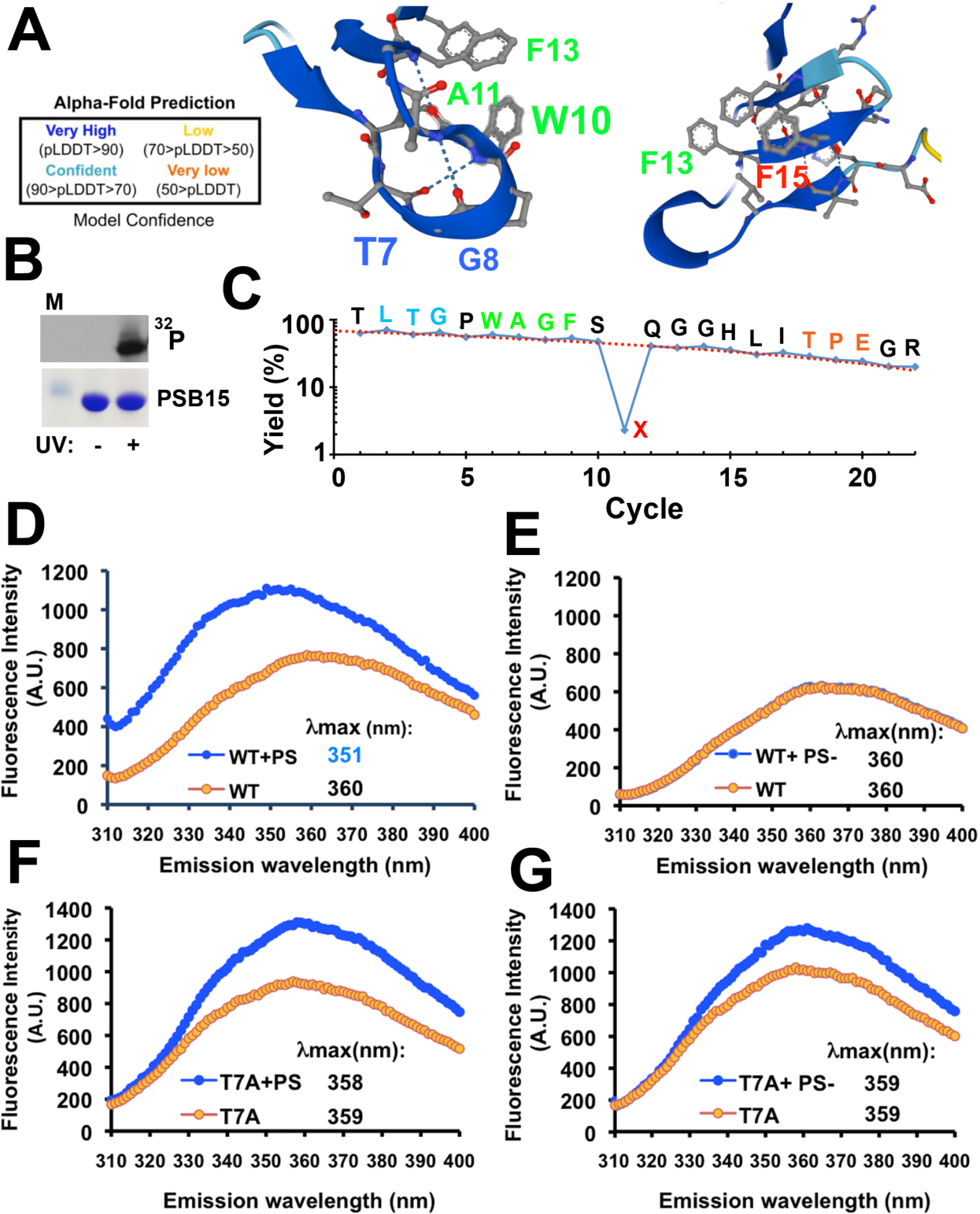
Identification and conformation changes of PSB15 DNA binding motif. (A) AlphaFold prediction of PSB15 N-terminus LTG (blue) and WAGF (green) motif. F15 is labeled in red. Model confidence is shown at left. (B) UV cross-linking of recombinant PSB15 and ^32^P-labeled PS DNA. Samples were analyzed in 15% SDS-PAGE (lower panel) and autoradiography (upper panel). (C) UV-crosslinked PSB15-PS complex was trypsinized, and ^32^P-labeled peptides were purified by HPLC then subjected to Edman sequencing. The recovery yield of phenylthiohydantoin (PTH)-amino acids is shown. (D-G) Intrinsic tryptophan fluorescence (W10) of PSB15 WT(2WA) (“WT”, D and E) and T7A(2WA) (“T7A”, F and G) upon PS and PS(-) DNA binding. Samples were excited at 295 nm, and fluorescence emission spectra was recorded between 310 to 400 nm.

To identify the direct DNA binding site, recombinant PSB15 protein was UV cross-linked with ^32^P-labeled PS DNA ([^32^P]PS probe). Followed by nuclease digestion, PSB15 was radiolabeled by the [^32^P]PS probe under UV irradiation (Figure 3B). After trypsin digestion, radiolabeled PSB15 peptides were separated by reverse phase HPLC and analyzed by Edman degradation sequencing (Merrill et al., 1984; Williams and Konigsberg, 1991; Prasad et al., 1993; Katsuki et l., 2005). A radiolabeled peptide with an amino acid sequence of T<u>LTG</u>P<u>WAGFS**X**</u>QGGHLI<u>TPE</u>GR, corresponding to amino acid 5 to 26 with LTG/WAGFXF/TPE motifs, was identified. Moreover, the recovery yield of phenylthiohydantoin (PTH)-phenylalanine was drastically reduced to 2.3% in cycle 11 (F15), indicating that F15 is the direct contact residue crosslinking to the PS DNA (Figure 3C). This result is also consistent with F15A abolishing PS binding (Figure 2E, 2F, and 2H).

Several hydrogen bonds within or between the ^6^LTG/^10^WAGFXF/^22^TPE motifs (e.g. G8-A11, W10-F13, W10-T7, F13-L6, T22-E24, T22-R26) were predicted by AlphaFold. These bonds appear to be critical to form the PSB15 ϕ hairpin (Figure 3A). We took advantage of tryptophan 10 (W10) being located within the immediate vicinity of the direct DNA binding site, F15 (**^10^W**AGFX**^15^F**) (Figure 3A). Tryptophan is most strongly influenced by the polarity of its local microenvironment, so it can serve as an unlabeled fluorescence sensor for detecting changes in its exposure to solvent (Ghisaidoobe and Chung, 2014; Fernández et al., 2016). This allows us to verify PS DNA binding properties of PSB15 using intrinsic tryptophan fluorescence. PSB15 contains three tryptophan residues (W10, W35, and W46). Two alanine substitutions at W35 and W46 in helix 1 of WT PSB15 protein [WT(2(WA)] did not affect DNA binding (Figure 2E and 2H, Table S2). Therefore, recombinant WT(2WA) protein was used to detect tryptophan fluorescence changes of the WAGFXF motif after binding to PS DNA. Following excitation at 295 nm, where the tyrosine absorption is negligible, the emission spectra of WT(2WA) showed a maximum (λ_max_) at 360 nm. λ_max_ was blue-shifted to 351 nm upon PS DNA binding, and fluorescence intensity significantly increased (Figure 3D). This data suggests that DNA binding caused the movement of W10 to the less polar inner core of the hydrophobic cavity of PSB15. This spectral change is specific to the PSB15-PS DNA interaction, because λ_max_ and intensity were neither changed with PS(-) DNA (λ_max_ =360 nm, Figure 2E) nor with F15A(2WA) protein (λ_max_ =358 nm, data not shown), in both conditions where no DNA binding occurred. Taken together, we conclude that PS DNA binding affects the tertiary structure of the ^10^WAGFXF motif.

Since T7 may interact with W10 via hydrogen bonding, we further tested our hypothesis that T7A alters PS binding specificity due to its influence with WAGFXF motif (Figure 3A). The T7A(2WA) protein has a comparable DNA binding affinity and specificity as T7A (Figure 2E, 2H, and Table S2). T7A(2WA) λ_max_ was 358 nm. In contrast to WT(2WA), there was no wavelength shift regardless of DNA binding (Figure 3F). Importantly, both PS and PS(-) DNA caused a similar increase of the W10 fluorescence of T7A(2WA), preserving the intensity gap observed between the curves in both figures (Figure 3F and 3G). This data suggests that T7 in the LTG motif contributes to the conformational change of the WAGFXF motif upon DNA interaction, which is essential to recognize the specific PS elements or exclude binding to PS(-).

### Basic helix 2 and C-terminal tail mediate PSB15 binding to the inner membrane and phospholipid cardiolipin

We next examined the subcellular localization of PSB15. Periplasmic, cytoplasmic, IM, and OM fractions from pPSB15-transformed Xcc-TcR cells were prepared. Ectopically expressed PSB15 was mostly detected in the IM fraction (Figure 4A). After infection with ϕLf-UK phage for 6 hours, the IM and OM of Xcc-TcR cells were separated by discontinuous sucrose density gradient centrifugation (Figure 4B) (Woodruff and Hancock, 1989; Doerrler et al., 2004). Viral PSB15 protein was cofractionated with the IM marker, MsbA, at the buoyant density of 1.14 g/ml, but not with the OM marker, OprF, at 1.22 g/ml. Both results confirmed that PSB15 is associated with the IM intracellularly.

**Figure 4.**
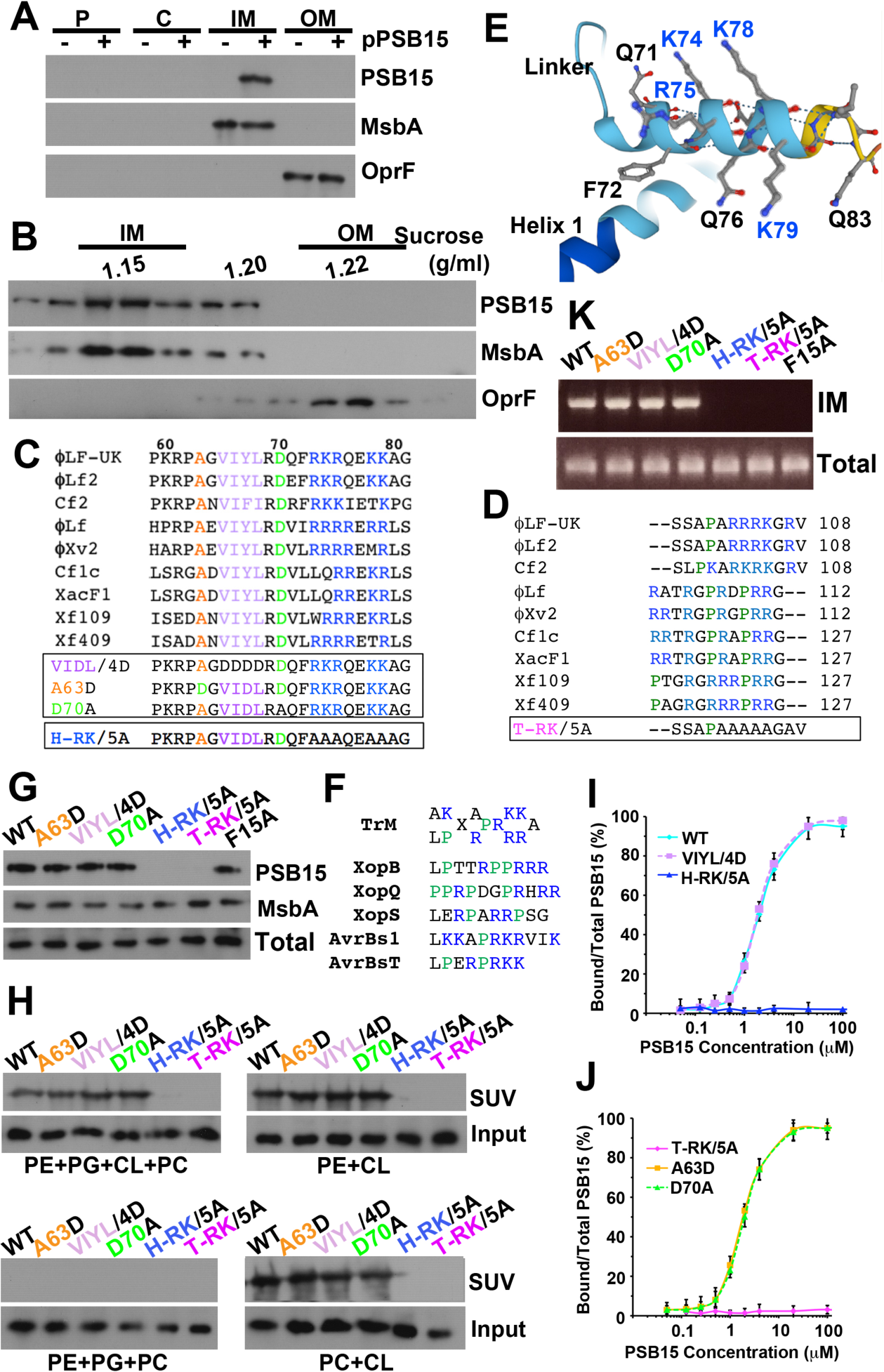
Membrane and phospholipid binding of PSB15. (A) Xcc-TcR bacteria was transformed with pPSB15 plasmid and fractionated as periplasmic (P), cytoplasmic (C), IM, and OM. Samples were analyzed in 15% SDS-PAGE and immunoblotting. MsbA and OprF are the IM and OM marker, respectively. (B) The IM and OM of <Lf-UK-infected bacteria was fractionated on a step sucrose gradient (25 to 61% [wt/wt]) centrifugation, and the fractions were analyzed by immunoblotting. Sucrose density was determined from the refractive index. (C and D) Alignment of PBS15 hydrophobic linker, basic helix 2 (C) and C-terminal tail (D) of *Xanthomonas* filamentous phages. Mutations are listed in the box. (E) AlphaFold prediction of PSB15 basic helix 2 structure. (F) Putative translocation motif (TrM) for *Xanthomonas* type III effector proteins (33). (G) pPSB15 mutant plasmids were transformed into Xcc-TcR bacteria. The IM was fractionated as (A) and immunoblotted. Total lysate (total) was probed with PSB15 as the control. (H) Liposome flotation assay. 5 μM PSB15 proteins were incubated with 1mM liposome (PE+PG+CL+PC=52%: 13%: 30%: 5%; PE+CL: 50%: 50%; PE+PG+PC: 40%: 40%: 20%). The mixtures were centrifuged in a step sucrose gradient (2M, 0.75 M, 0 M). The floated liposome-bound (SUV) fractions were collected and analyzed by SDS-PAGE and immunoblotting. Inputs were loaded as the control. (I and J) Binding kinetics of GFP-PSB15 and FM4-64-labeled liposome (PE: PG: CL: PC=52%: 13%: 30%: 5%). Flotation assays were carried out as (H). Fluorescence intensity of GFP-PSB15 was measured and normalized with FM4-64 intensity. (K) Xcc-TcR was transformed with pPSB15 mutant plasmids as indicated, and bacteria were infected with ϕLf-UK-ϕPSB15 phage. The IM was fractionated and cross-linked by formaldehyde, and phage DNA was immunoprecipitated using anti-PSB15 antibody. Precipitated phage DNA was recovered and amplified by PCR (nt 190 to 476 of ϕLf-UK) and analyzed in 1% agarose gel. DNA from cytoplasmic fraction was PCR amplified by the same primers and served as the control (total).

PSB15 contains a hydrophobic linker (A55-L68), Arg/Lys-rich helix 2 (R69-G81), and Pro/Arg/Lys-rich C-terminal tail (AA L82-V108) (Figure 4C, 4D, 4E, and 5A). Because both the helix 2 and C-terminal tail are very basic, the isoelectric points of PSB15 proteins are estimated within the range of 10.47 to 11.35. The prominent feature of Pro/Arg/Lys-rich sequences is also found in the putative translocation motif (TrM) for *Xanthomonas* type III effector proteins (e.g. XopB, Figure 4F), which is required for their association with bacterial membranes (Prochaska et al., 2018). To investigate the membrane binding properties of PSB15, we generated substitution mutations of conserved residues in the linker (A63D; ^65^VIYL → DDDD, named “VIYL/4D”), helix 2 (D70A; ^73^RKRQEKK→ AAAQEAA, named “H-RK/5A”), and C-terminal tail (^102^RRRKGRV→ AAAAGAV, named “T-RK/5A”) in pPBS15 plasmid (Figure 4C and 4D). Ectopically expressed linker (A63D and VIYL/4D), D70A, and F15A proteins can be detected in the IM fraction (Figure 4G). However, loss of basic residues (H-RK/5A and T-RK/5A) resulted in the failure of PSB15 targeting to the IM. Our results indicate that the basic helix 2 and the C-terminal tail are PSB15 membrane binding motifs.

Our data also led to the next hypothesis that PSB15 helix 2 and the C-terminal tail interact with the negatively charged membrane surface. It has been shown that *Xanthomonas* membranes are rich in an anionic phospholipid, cardiolipin (CL), which carries two negative charges at physiological pH (Moser et al., 2014; Prochaska et al., 2018). The direct binding of PSB15 with phospholipids was tested by mixing recombinant PSB15 with liposomes (small unilamellar vesicles, SUVs) in different combinations of phospholipids (phosphatidylethanolamine, PE; phosphocholine, PC; phosphatidylglycerol, PG; and CL). Liposome flotation assay was used to separate SUV-bound and free PSB15 by a step sucrose gradient. Figure 4H shows that PSB15 was bound to SUVs (upper) with composition similar to the native *Xanthomonas* membrane (PE: PG: CL: PC=52%: 13%: 30%: 5%, “native” SUVs) (Moser et al., 2014; Prochaska et al., 2018). PSB15 can bind to liposomes with CL (PE: CL= 50%: 50% and PC: CL= 50%: 50%), but not with negatively charged PG (PE: PG: PC=40%: 40%: 20%), indicating that CL is the lipid-binding target. Binding affinity between PSB15 and native SUVs was estimated as K_d,app_ 2.3 ± 0.3 μM. A similar binding affinity of GFP-PSB15 and FM4-64-labled native SUVs, measured by fluorescence, was estimated with K_d,app_ (2.5 ± 0.4 μM, Figure 4I and 4J, Supplemental Table S3). Mutations of the linker (A63D, VIYL/4D), D70A or the N-terminus did not affect SUV binding, but both H-RK/5A and T-RK/5A lost their lipid-binding activity (Figure 4I and 4J, Table S3). We concluded that Arg/Lys residues in the helix 2 and C-terminal tail are essential to target PSB15 to the membrane domain by binding to CL.

### PSB15 targets phage DNA to the IM

Because PSB15 binds to both PS DNA and the IM at different domains, we tested the hypothesis that PSB15 recruits phage DNA to the membrane. pPSB15 mutant plasmids were transformed into bacteria, followed by ϕLf-UK-ϕPSB15 phage infection. The IM was fractionated and collected, followed by formaldehyde cross-linking and chromatin immunoprecipitation using anti-PSB15 antibody. After PCR amplification, the antibody-precipitated phage DNA was able to be detected in PSB15 WT, A63D, VIYL/4D, and D70A, but not in H-RK/5A-, T-RK/5A-, and F15A-expressing cells. This result indicates that lipid- and DNA-binding domains are required for PSB15 targeting phage DNA to the IM. Moreover, mutants in each domain failed to rescue assembly arrest of ϕLf-UK-ϕPSB15 (Table S2). We concluded that IM targeting of PSB15 and PS DNA is an essential checkpoint for phage assembly.

### Thioredoxin binds to and releases PSB15-PS DNA complex from the lipid membrane

Thioredoxin (Trx) is the only known host protein essential for Ff assembly to date (Russel and Model, 1985; Lim et al., 1985). Without Trx, coat protein accumulates in the IM, but particle extrusion from the cell is abolished (Russel, M. and Model, 1983). However, Trx’s function in phage assembly has not been well defined since then. Russel and Model showed genetic evidence that *trxA* mutations of *E. coli* (Pro34, Gly74, Pro76, Gly92) can still participate in redox reactions but did not support Ff phage assembly. They also suggested that Trx can interact with thioredoxin reductase and phage assembly machinery dually on the same surface site (Russel and Model, 1986). We found that Xcc-TcR bacteria with TrxA G93D mutation also failed to support ϕLF-UK phage assembly (Figure 1C). The essential role of both PSB15 and Trx in phage packaging prompted us to examine their interaction. We expressed recombinant GST-tagged *Xanthomonas campestris* TrxA and its G93D mutant (mimic to Ff assembly mutant G92D) in *E. coli* and purified them in a glutathione affinity Sepharose column. After mixing purified TrxA and PSB15, the protein complex was pulled down using glutathione resin. WT PSB15 can be co-pulled down with GST-TrxA, but not with GST-TrxA G93D mutant (white arrowhead, Figure 5B), indicating that *Xanthomonas* TrxA G93 residue is critical for its direct binding to PSB15. We further mapped the TrxA binding site(s) of PSB15. A63D, VIYL/4D and D70A mutants failed to bind to TrxA, while the TrxA binding ability of R69A, DNA-binding (LTG/3A, WAGF/4A, TPE/3A, F15A) and lipid-binding (H-RK/5A, T-RK/5A) mutants were comparable to WT. This data indicates that the hydrophobic linker and D70 are essential for TrxA binding, whose interaction is also essential to support phage assembly (Table S2).

**Figure 5.**
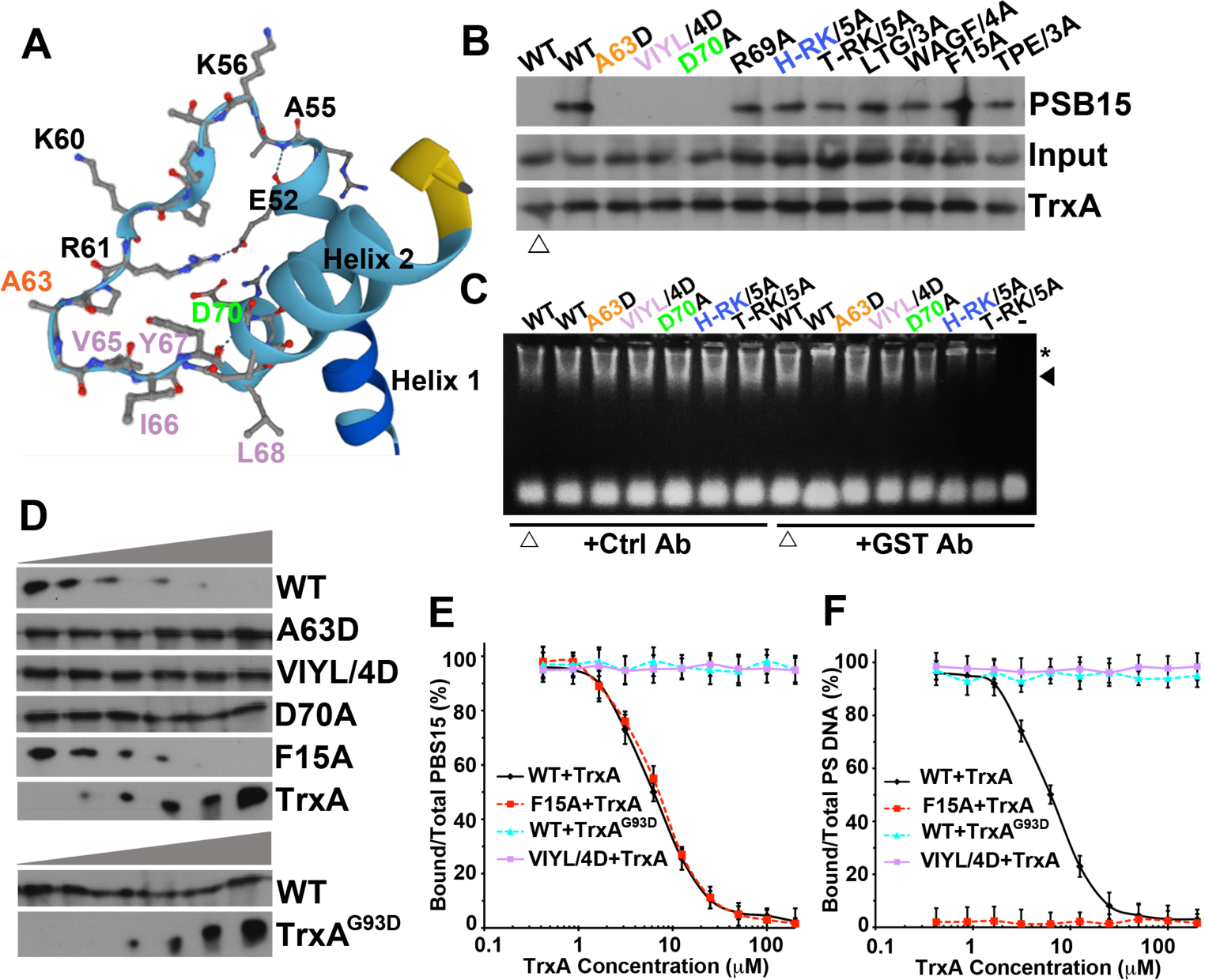
Trx binding and its effects on PSB15. (A) AlphaFold prediction of PSB15 hydrophobic linker and helix 2. (B) Pull down assay of GST*-*TrxA with PSB15. TrxA mutant G93D was used as the negative control (white arrowhead). (C) TrxA-PSB15-PS complex (black arrowhead) was analyzed in EMSA. The supershift by anti-GST antibody (asterisk) was shown. Lanes with TrxA-G93D were labeled with white arrowhead. (D) Thioredoxin was added with PSB15 and liposome (PE:PG:CL:PC= 52: 13: 30: 5) for 30 min. Mixtures were subjected to a floatation assay. Liposome-bound PSB15 and input of TrxA (and G93D mutant) were analyzed by immunoblotting. (E and F) Kinetics of liposome-bound GFP-PSB15 (E) and FITC-labeled PS DNA (F) in the flotation assay in response to TrxA and TrxA G93D concentration. F15A acts as a negative control, which confirmed that PS DNA is bound to liposomes via PSB15.

TrxA-binding mutants A63D, VIYL/4D, and D70A can still bind to PS DNA (Table S2) and the lipid membrane (Figure 4G-4J). Therefore PS DNA-, membrane-, and TrxA-binding are at different PSB15 domains. We next tested if PSB15 can interact with TrxA and PS DNA simultaneously. GST-TrxA can form a complex with PS DNA and PSB15-WT or lipid-binding mutants (H-RK/5A and T-RK/5A), as evident by a supershift band caused by anti-GST antibody (Figure 5C). In contrast, anti-GST antibody did not retard the PSB15-PS DNA complex with GST-TrxA G93D or PSB15 TrxA-binding mutants (A63D, VIYL/4D, D70A). Negative control shows that neither TrxA nor anti-GST antibody alone was able to form a complex with PS DNA, confirming that the supershift is dependent on TrxA binding to PSB15. Moreover, TrxA binding did not affect PSB15-PS binding affinity (K_d,app_ 12.2 ± 1.3 nM). Taken together, our data supported that TrxA and PS DNA can bind to different domains of PSB15 as a complex.

Because TrxA-binding sites of PSB15 (A63-D70) are adjacent to the lipid-binding domain (R73-K79 in basic helix 2) (Figure 4C and 5A), it seems reasonable to hypothesize that TrxA binding affects PSB15 association with the lipid membrane. This guided the investigation of their interaction. Recombinant TrxA was mixed with PSB15 and native SUVs in a flotation assay. TrxA was not detected with SUVs, indicating that neither lipid nor PSB15 recruited TrxA to the membrane (data not shown). However, SUVs-bound WT PSB15 or F15A levels were reduced as TrxA input concentration increased (Figure 5D). Membrane-bound PSB15 was unchanged when incubated with TrxA binding mutants (A63D, VIYL/4D, D70A) or TrxA G93D. This indicates that TrxA binding releases PSB15 from SUVs. The fluorescence intensity of recombinant GFP-PSB15 was also reduced in association with FM4-64-labeled native SUVs at increasing concentration of TrxA (Figure 5E).

The possibility that the PSB15-PS DNA complex can also be released together by TrxA from liposomes was also tested. Figure 5F shows a concentration response curve for SUV-bound, FITC-labeled PS DNA decreasing in the presence of TrxA, but not TrxA G93D or PSB15 VIYL/4D. In addition, the response curves show no difference when TrxA was added simultaneously with or after PSB15-PS bound to SUVs (data not shown). Taken together, these findings support the idea that TrxA dissociates the PSB15-PS DNA complex from the phospholipid membrane *in vitro*.

### Trx and DNA binding facilitate PSB15 dissociation *in vivo*

Our results led to the hypothesis that Trx can regulate PSB15 dynamics at the IM. To date, there is very little literature about the direct observation of filamentous phage proteins in bacteria using live cell imaging. In order to explore *in vivo* behavior of PSB15, we visualized ϕLf-UK-GFP-PSB15 phage-infected cells using total internal reflection fluorescence (TIRF) microscopy, which typically only penetrates samples up to 100 nm, and minimizes photobleaching (Axelrod, 2001; Johnson and Vert, 2017). Live cell imaging showed that GFP-PSB15 was temporarily distributed only at the cell pole membrane as puncta (Figure 6A). Although the fluorescence intensity of GFP-PSB15 puncta fluctuated after it reached a plateau, the dwell time of each punctum from multiple cells can be combined to produce a robust mean value (Figure 6A, movies S1). The mean dwell time was calculated to be approximately 8.23 ± 1.30 s from WT PSB15 puncta (Figure 6B). We then tested whether TrxA existence or binding could affect PSB15 behavior *in vivo*. Either loss of TrxA binding activity (A63D) or in Xcc-TcR TrxA G93D bacteria did not interfere with the localization of PSB15 at the poles (Figure 6A). However, the mean dwell time of PSB15 puncta was more than 50 s in both conditions (Figure 6A and 6C, movies S2 and S3).

**Figure 6.**
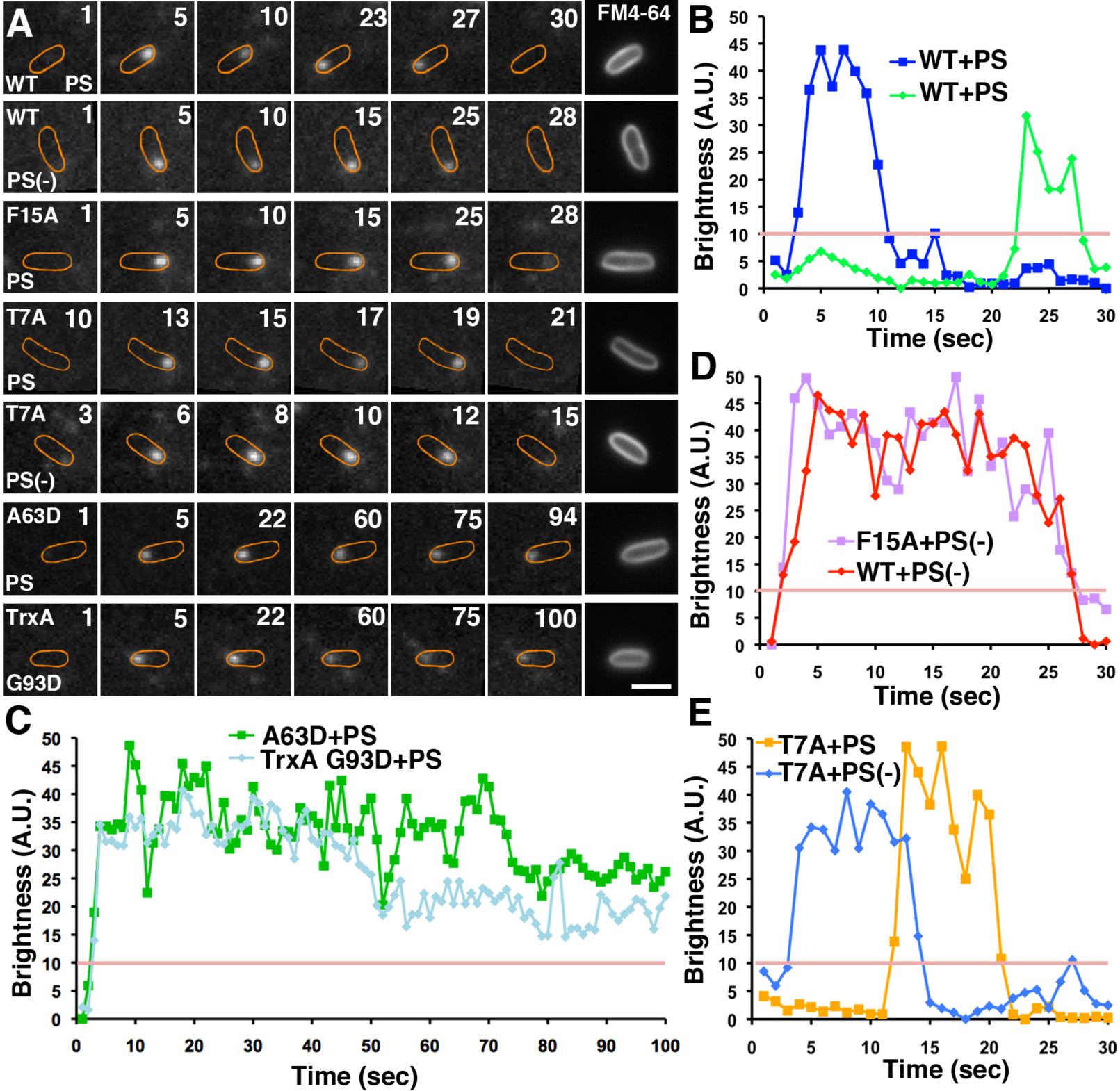
PSB15 dynamics in *vivo*. (A) Live cell imaging of GFP-PSB15 under TIRF microscopy. pGII-transformed Xcc-TcR bacteria were infected with <Lf-UK GFP-PSB15 or PS(-)-GFP-PSB15 phages at a multiplicity of infections of 1 for 30 min. Snapshots of time-lapse movies were shown. The timestamps are labeled on the upper right corner (sec) in the corresponding movies (Movies S1-S7) and the IM was labeled by FM4-64 dye and margined. Bar = 1 μm. (B-E) Example of brightness of GFP-PSB15 spots at cell membrane was recorded in (A). Dwelling time is defined as the duration of GFP signal more than 10 arbitrary unit (A.U.; pink line)

Although the localization and intensity of PSB15-F15A was unchanged in infected bacteria, the mean dwell time increased significantly to 25.6 ± 3.3 s (*p*< 0.001, Mann-Whitney *U* test). Similar results were visualized with PSB15 in the presence of PS(-)-GFP-PSB15 phage (mean dwell time=26.3 ± 4.1s; *p*< 0.001, Mann-Whitney *U* test) (Figure 6A and 6D, movies S4 and S5). These findings prompted the hypothesis that DNA binding activity facilitates either the release of PSB15 from the pole membrane or the dissociation of punctate structure *in vivo*. We further tested this hypothesis by using ϕLf-UK-GFP-PSB15-T7A or PS(-)-GFP-PSB15-T7A phages, where T7A was shown to bind to PS and PS(-) DNA at indistinguishable kinetics (Table S2). T7A punctae showed a similar dwelling time (9.23 ± 1.81 s and 8.98 ± 1.60 s, respectively) between both PS and PS(-), and both were comparable to PSB15-WT with PS (Figure 6A and 6E, movies S6 and S7). Taken together, our findings indicate that both Trx and DNA binding facilitate PSB15 dissociation from the pole membrane *in vivo*.

### Discussions

Encapsidation of asymmetric filamentous phage particles is a highly ordered reaction, which starts with the targeting of viral ssDNAs to the IM, where the coat proteins are located (Wickner and Killick, 1977; Russel, 1991). Our findings provide the first evidence of a phage assembly checkpoint that ensures membrane conditions are favorable for phage assembly to proceed. This checkpoint is controlled by a nonstructural phage protein, PSB15. PSB15 targets PS DNA to the IM for efficient assembly *in trans*, which has not been reported in coliphages or other filamentous phages. Previously, Hasse et al. reported that the M13 pV-phage DNA complex, but not pV alone, is targeted to pI/pXI in the liposome *in vitro* (Hasse et al. 2022). The Ff PS appears to load onto pI/pXI machinery, where it uses the platform to bind to pVII/pIX (Russel 1991; Dotto and Zinder, 1991; Grant and Webster, 1984a; Grant, R.A. and Webster, 1984b). We found that the integrative filamentous phages PS plays a more active roles in IM targeting/dissociation via PSB15 and Trx. It serves as a critical checkpoint that most likely occurs prior to the addition of coat proteins. Failure of this accurate virion formation quality control mechanism causes the arrest of assembly reactions.

Our current working model of the phage assembly checkpoint is illustrated in Figure 7. PSB15 increases the concentration of viral ssDNAs by targeting them to the pole of the IM (the “priming” site), thus facilitating virion production by increasing the rates of intermolecular assembly reactions. Viral coat proteins contain information essential for specific assembly, but accurate encapsidation usually also requires scaffolding or chaperon proteins from the hosts or the viruses themselves (Palucha et al., 2005; Dearborn et al. 2017; Mateu 2013). PSB15 targeting the phage DNA to the IM is a highly active checkpoint enabling loading of the priming site. Other protein factors may also restrict or regulate the number of interactions in which viral nucleic acids and coat proteins are involved, such as in the formation of the “initiation complex” proposed previously (Feng et al., 1999). Once this checkpoint is satisfied, PSB15-phage DNA is released from the priming site by Trx, and phage DNA is loaded on the pI export complex (Haase et al., 2022). Finally, other coat proteins (pVII, pVIII and pIII/pVI) are added during egress to generate mature viral particles. The possibility of PSB15 and thioredoxin being involved in (1) coat protein transitions from their IM-embedded to structural virion forms, and (2) the release of phage particles, still requires further investigation.

**Figure 7.**
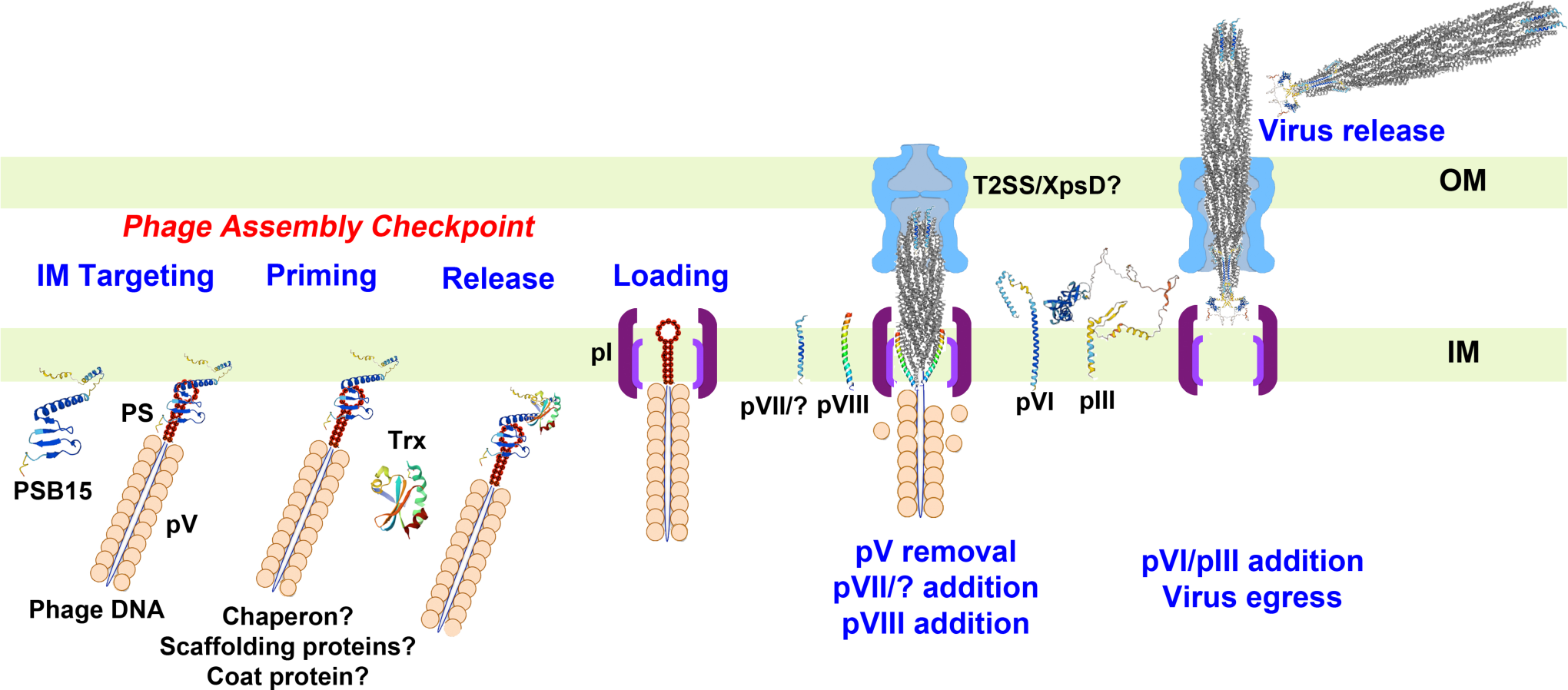
Working model of filamentous phage assembly and its checkpoint control. PSB15 efficiently targets phage DNA to the IM through PS binding at the priming site in the cell pole. Other essential factors (chaperons, scaffolding factors etc.) may participate in the prerequisite reactions and/or the formation of the “initiation complex” to satisfy the assembly checkpoint. Once the checkpoint is satisfied, Trx releases PSB15 and phage DNA from the priming site, then phage DNA is loaded to pI export complex. pV is replaced by major coat protein pVIII, driven by ATP hydrolysis of pI. Virus particles assemble and pass through the OM (likely via the host secretin complex, such as type II secretion system T2SS). pVI and pIII are added to terminate the assembly process, and virions are released out of the host. The structures were illustrated based on AlphaFold or PDB as the following: PSB15 (Uniport: Q8P905), pIII (Uniport: Q3BSR6), pVI (Uniport: Q3BSR8), pVII (Uniport: Q3BSR4), pVIII (PDB:2IFO), thioredoxin (PDB: 2TRX), phage (PDB: 2IFO, 8B3P, 8B3O, 8IXK, 8IXL). Both ends of phage are adopted from f1 and M13 phage structures (Conners et al., 2023; Jia and Xiang, 2023).

Both Ff and *Xanthomonas* phage PS are DNA harpin structures, and their mutations cause tiny plaques to form at high frequencies, generating pseudorevertants with near normal sized plaques (Russel and Model, 1989; Yeh et al., 2023). It is worthy to note that the PS(-) sequence still maintains the consensus stem sequence (GGCACCGCG), but contains T340G/C344G mutations in the loop. PSB15 T7A and T22A can bind to PS(-) as interaction suppressors, and T7A can rescue PSB15 dwelling time to the WT dwelling time. Therefore, we speculate that T7A and T22A mutants appear to relieve the stringency of binding specificity and packaging constraints at the loop of T340 and C344. AlphaFold prediction suggested that T7 and W10 interact with each other via a hydrogen bond (Figure 3A). Our intrinsic tryptophan fluorescence data also reflected that the hydrophobicity and microenvironment of W10 residue is affected by PS DNA binding. T7 may participate in DNA binding specificity via a change of the WAGFXF motif conformation.

It has been shown that the XafT protein of the *Vibrio* TLCΦ phage, with both helix-turn-helix XRE and DUF3653 domains, is involved in XerC/D-mediated phage DNA integration by interacting with and activating XerD (Midonet et al., 2019; Miele et al., 2022). Because the helix-turn-helix XRE is a DNA binding domain, whether the XafT C-terminal DUF3653 domain directly binds to DNA remains unclear. In *Xanthomonas* and *Stenotrophomonas* phages, XRE HTH and DUF3653 are encoded by two independent ORFs in the negative strand. The DUF3653 domain is located at the N-terminus of PSB15. In contrat to XafT, loss of PSB15 does not affect phage DNA integration efficiency, indicating that PSB15 and its DUF3653 domain are nonessential for XerC/D recombination. On the other hand, we confirmed that PS DNA binds directly at F15 in the DUF3653 domain. Since our attempt to search for PSB15 DNA binding sites via the mass spectrometry method was unsuccessful, other possible DNA binding site(s) cannot be completely ruled out yet. Whether other hundreds of DUF3653 domain-containing proteins from 51 bacterial species are involved in either XerC/D recombination or phage assembly needs further investigation.

TrxA function and its binding protein(s) in Ff phage assembly have not been well studied in 40 years (Russel and Model, 1983; Russel and Model, 1985; Russel and Model, 1986; Lim et al., 1985). It has been proposed that Trx may confer processivity by stripping pV off of the DNA and replacing it with the major coat protein, pVIII (Russel, 1991). Ff gI missense mutations can grow on *E. coli* hosts containing certain *trx*A missense mutations, suggesting that pI and Trx genetically interact. However, their physical binding has never been documented (Russel and Model, 1983). Here we reported the first evidence that PSB15 can bind to TrxA, working together as a control assembly checkpoint. Trx binding is not required for the targeting of PSB15 to the IM or cell poles, but it is essential for the timely release of PSB15 from the membrane *in vitro* and *in vivo*. Both the C-terminus and the basic helix 2 residues are required for binding to the negatively charged cardiolipin of the IM, likely via electrostatic interactions. The isoelectric point value of *Xanthomonas* TrxA is 5.17. Since TrxA’s binding site is adjacent to basic helix 2 (Figure 4C and 5A), TrxA binding may neutralize the positive charges of basic helix 2 and interfere with the PSB15-cardiolipin interaction. Given the fact that both PS- and Trx-binding activity facilitate membrane release of PSB15 *in vivo*, identification of PSB15- and Trx-binding protein(s) will help us understand the mechanism of assembly checkpoint control.

Along with PSB15: gVII, gI, and ORF11 genetically interact with the PS. The characterization of gVII, gI, and ORF11 suppressors and their functions are currently underway. It is worthy to note that mutations to either the PS or these assembly proteins in Ff and *Xanthomonas* phages significantly reduce, but do not completely terminate, viral assembly. Whether or not these virions are packaged via an alternative pathway (e.g. pV targets the phage DNA directly to pI/pXI) requires further exploration (Haase et al., 2022).

## METHODS

### Bacteria and plasmids

The host bacteria, Xcc-TcR, was maintained by tetracycline as described (Yeh et al., 2023). TrxA G93D mutation was introduced in Xcc-TcR by homologous recombination in which TrxA G93D was expressed in *cis* under control of the native promoter (Kaniga et al., 1991; Huguet et al., 1998).

The ϕLf-UK replicative form (RF) DNA, the karamycin-resistant pORIPS plasmid with ϕLf-UK *ori* and PS sequence (245-365, MH206184), as well as chloramphenicol-resistant pGII plasmid was maintained as previously (Yeh et al., 2023). A stop codon was introduced into the start codon of ϕLf-UK ORF9 gene (nt 4983-5309, MH206184) of RF DNA to generate ϕLf-UK-ϕPSB15. The T340A/C341G mutation, which results in the synonymous mutation of serine 114 of the gII protein but losing PS activity, was introduced in ϕLf-UK and ϕLf-UK-ϕPSB15 RF DNA to generate ϕLf-UK T340A/C341G and T340A/C341G-ϕPSB15. The antisense PS sequence [PS(-), 5’-CACGGGTCCGGTTGTAGAGGCCCGCG-3’] was introduced to replace the PS sequence of ϕLf-UK-ϕPSB15 RF DNA to generate PS(-)-ϕPSB15. The ORF9 gene of ϕLf-UK was PCR amplified and subcloned into a broad-host-range expression, chloramphenicol resistance vector, pHP11, as pPSB15 plasmid (Reece and Phillips, 1995).

To obtain the constructed pR-PSB15 expressing recombinant PSB15 protein, the ϕLf-UK ORF9 gene was PCR amplified with an extra enterokinase cleavage site (DDDDK) upstream of the starting codon. The PCR fragment was subcloned in the *Bam*HI and *Not*I site of pET-28a+GFP-CAMSAP1 CKK (Addgene #59033), which contains 6 histidine (His) and superfolder GFP (sfGFP) at its N-terminus (Hendershott and Vale, 2014). These constructs were transformed and selected by either kanamycin (30µg/ml) or chloramphenicol (200 µg/ml). The ORF9 gene of ϕLf-UK RF DNA was replaced with the coding region of sfGFP-PSB15 of pR-PSB15 to generate ϕLf-UK-GFP-PSB15. The PS sequence was replaced by the PS(-) to generate PS(-)-GFP-PSB15. The *Xanthomonas* thioredoxin A (TrxA, NZ_CP058243) sequence was synthesized by GeneScript and subcloned into the pDEST15 vector (ThermoFisher Scientific) to create pDEST-GST-TrxA. TrxA G93D mutation was introduced to generate pDEST-GST-TrxA-G93D. Plasmids were purified by Plasmid Maxi Kit according to Qiagen’s protocol. Site directed mutagenesis was carried out using the QuikChange II Site-directed mutagenesis kit (Agilent).

### Molecular Biology and detection of phage particles

Plasmid DNA propagation and electroporation was described previously^24,55,56^. RF DNAs of ϕLf-UK-ι1PSB15 and PS(-)-ι1PSB15 mutants were co-transformed with pPSB15 and pGII in Xcc-TcR to measure their phage production yield. Purification of phage particles, as well as phage DNA, RF DNA, and the host chromosomal DNA were described, respectively (Yeh, 2017; Yeh 2020). Phage integration was analyzed by PCR using oligonucleotide primers for ϕLf-UK. Quantitative PCR (qPCR) analysis of phages released in medium were measured based on our published methods (Yeh et al., 2023). PSB15 protein sequences were aligned by Clustal Omega, and the structure was predicted by AlphaFold.

### Genetic screening of suppressor mutations for ϕLf-UK PS mutant phage

Russel and Model’s method was adopted for the first PS suppressor screening (Russel and Model, 1989). ϕLf-UK T340A/ C341G RF was transformed into Xcc-TcR. 0.3 ml of the exponentially growing culture of Xcc-TcR (OD_600_=0.6) was mixed with 40 μl of RF-transformed cells. The mixture was plated together with 3 ml of soft agar on a LB plate and then incubated at 28°C for 18 hours. One clone with tiny morphology was chossen, and the soft agar plating was repeated. Normal-sized plaques (>1 mm diameter) were selected under an Olympus SZH dissecting microscope, and the phages were amplified. For the second screening, these helper phages were used to infect the Xcc-TcR cell transformed with pGII and pORIPS-PS(-) plasmid, which contains the PS(-) sequence. The copy numbers of transducing particles (TPs) released in the medium were measured by qPCR (Yeh et al., 2024). The RF DNAs of pseudorevertants (with normal TP release) were isolated, and the whole phage genome sequences were analyzed by Sanger sequencing. A total of 4 independent suppressor screens were performed, and 25 pseudorevertant genomes were completely sequenced for each screen by PCR.

### Purification and analytical size-exclusion chromatography of PSB15 proteins

Recombinant ϕLf-UK PSB15 protein was expressed in B834 (DE3) pLysS cells (EMD Millipore) by transforming pR-PSB15. Bacteria were grown in LB medium at 37 °C to A600 0.8, supplemented with 0.1 mM isopropyl β-D-1-thiogalactopyranoside, and then grown at 18 °C for an additional 18 hours. 50 ml of bacterial culture was harvested and resuspended in 10 ml BugBuster solution (EMD Millipore) with 1% Triton X-100, 0.5% CHAPS, 1 mg/ml lysozyme, 5 U/ml Benzonase, protease inhibitors, and 2 mM β-mercaptoethanol (BME), and then incubated at room temperature for 1 hour. The later steps were performed at 4 °C. The bacterial lysate was clarified by 30-min centrifugation (40,000 *g*). The inclusion bodies in the pellets were suspended with 10 ml Tris-urea buffer (20 mM Tris pH 7.9, 150 mM NaCl, 6M urea) overnight. Polyethyleneimine (PEI) solution (Sigma Aldrich 40872-7) was added to give a final PEI concentration of 0.1 mg/mL (Nian et al., 2008). After vortexing and incubating for 30 min, DNA contamination and undissolved debris were removed by centrifugation at 40,000 *g* for 30 min. The supernatants were applied to a 2 ml nickel agarose (high density, GoldBio) for binding for 1 hour, and the resin was washed with 10 ml Tris-urea buffer followed by 10 ml Tris-urea buffer containing 25 mM imidazole. The proteins were eluted with 10 ml Tris-urea buffer containing 0.5 M imidazole. Fractions with PSB15 proteins were pooled and diluted with Tris buffer (20 mM Tris pH 7.9, 150 mM NaCl) in a 1:3 volume ratio for recombinant enterokinase digestion (Novagen, 1 unit/ 50 μg protein) overnight (Gasparian et al., 2006). Protease was removed according to the manufactural protocol (Novagen). PSB15 proteins were denatured by dialysis against one liter of CAPS [3-(cyclohexylamino)-1-propanesulfonic acid] buffer (20 mM CAPS, pH 11.0; 0.1 M NaCl; 1 mM BME; 10% glycerol) with 6M urea. After passing samples through the nickel agarose to remove the His-sfGFP tag, untagged PSB15 in the flow through was renatured with a gradual decrease of urea concentration (3 M, 1.5 M and 0 M) in a dialysis bag (Tsai et al., 1999). The refolded proteins were concentrated with Amicon 3 kDa Ultracel filters (EMD Millipore). For GFP-PSB15 purification, the steps of enterokinase digestion and His-sfGFP removal were omitted.

Analytical size-exclusion chromatography was performed using an AKTA fast protein liquid chromatography (FPLC) system (Amersham GE Healthcare). The concentrated samples were loaded into a Superose12 HR 10/30 FPLC column (GE) in CAPS buffer. Elution was performed at a constant flow rate of 0.4 ml/min, and the absorbance of the eluate was monitored at 280 nm. The molecular mass of the PSB15 protein was estimated from a calibration curve constructed from analysis of the following standards: insulin (5.7 kDa), chicken egg white lysozyme (14.3 kDa), and carbonic anhydrase (29 kDa). The peak fractions containing PSB15 proteins were verified by sodium dodecyl sulfate-polyacrylamide gel electrophoresis (SDS-PAGE) and Coomassie blue staining.

### Purification of GST-TrxA protein and binding to PSB15

Protein expression and purification of GST-TrxA and G93D mutant in *E. coli* using GSTtrap FPLC column (GE) was conducted as described previously (Yeh, 2020). Purified protein was dialyzed with protein binding (PB) buffer (50 mM Tris– HCl, pH 8.0, 100 mM NaCl, and 1 mM DTT) overnight at 4 °C, concentrated with Amicon 3 kDa Ultracel ultra centrifugal filters, and stored at -80 °C until being used.

10 μg GST-TrxA and PSB15 were mixed in 100 μl protein binding (PB) buffer at 4 °C for 1 hour. 20 μl PB buffer-washed glutathione agarose resin (1:1 slurry, ThermoFisher Scientific 16101) and a 380 μl PB buffer was added and incubated at 4 °C for 1 hour. After the resin was washed with 600 μl PB buffer with 0.1% Tween-20 for 5 times, the bound protein was eluted with 20 μl PB buffer with 10 mM glutathione at room temperature for 20 min. Eluted proteins were analyzed by 12% SDS-PAGE and immunoblotting.

### Antisera production of PSB15 and immunoblotting

Custom rabbit PSB15 antisera were prepared by ThermoFisher Scientific. The PSB15 antibody was affinity purified using CNBr-activated agarose. Protein samples were separated in SDS-PAGE and transferred onto the Immobilon-P PVDF membrane (0.45-µm-pore-size, EMD Millipore). PSB15 protein was detected using affinity purified antibody (1:2000) followed by alkaline phosphatase-conjugated anti-rabbit secondary antibodies (1:5000) and CDP-Star (0.25 mM) according to the manufacturer’s instructions (Roche).

### Identification of the DNA-binding residue of PSB15

Our method that identified the PSB15 DNA-binding site using Edman sequencing was modified from previous published protocols (Merrill et al., 1984; Williams and Konigsberg, 1991; Prasad et al., 1993; Katsuki et l., 2005). The PS oligonucleotide (5’-CGCGGGCCTCTACAACCGGCACCGTG) was synthesized by IDT and 5’-labeled with [γ-^32^P]ATP and T4 polynucleotide kinase (NEB) to generate the [^32^P]PS probe. 3 nmol (41.4 μg) of PSB15 protein and the [^32^P]PS probe were mixed together in an equal molar ratio in 15 μl buffer (25 mM Tris, pH 8.0, 10mM NaCl, 1 mM EDTA) at room temperature for 20 mins. The samples were spotted on parafilm paper and irradiated in a Stratagene UV Stratalinker 1800 Crosslinker (254 nm, 20 mJ/cm^2^) for 2 mins. The mixtures were precipitated with 10% ice-cold trichloroacetic acid, washed twice with acetone, and air-dried. The pellet was resuspended in 100 μl of 8 M urea. One tenth of cross-linked sample was analyzed in 15% SDS-PAGE and autoradiography. 9 out of the 10 of the samples were diluted 1:8 with 0.1 M NH_4_HCO_3_, pH 8.0., then 2 μg trypsin was added and the reaction mixture was then incubated at 37°C for 20 hrs. Cross-linked tryptic peptides were purified by a Vydac 218TP C18, 300 Å, 5 µm, 4.6 x 250 mm HPLC Column in Thermo Finnigan Surveyor HPLC. The peptides were fractionated every 30 s. 200 μl of each fraction was added to 2 ml of Ultima Gold liquid scintillation cocktail (Perkin Elmer), and the radioactivity was measured by a LS-6500 scintillation counter (Beckman). The fraction with the highest radioactivity was collected and pooled, concentrated, then subjected to Edman sequencing in a Shimadzu PPSQ-51A protein sequencer. The chromatogram was analyzed by PPSQ Postrun Analysis software (Shimadzu).

### Intrinsic tryptophan fluorescence measurements

PSB15 protein samples (40 μg) were mixed with 100 μM PS or PS (-) oligonucleotides in 200 μl buffer, pH 7.9, of 50 mM Tris-HCl, 1 mM MgCl_2_. After the samples were incubated in Greiner UV-Star 96 well plates (Greiner BIO-ONE 655801) for 30 min at 37°C, the fluorescence spectra were recorded on a SpectraMax Gemini EM spectrophotometer (Molecular Devices). The excitation wavelength of the intrinsic tryptophan fluorescence was 295 nm with medium PMT sensitivity. Tryptophan emission spectra were recorded from 310 to 400 nm in 1-nm increments. The fluorescence intensity was corrected for the inner filter effect by using the absorption values at the excitation and emission wavelength as described (Fernández et al., 2016; van de Weert, 2010). Each experiment was triplicated with three independent samples.

### Liposome preparation

1,2-dioleoyl-*sn*-glycero-3-phosphoethanolamine (PE), 1,2-dioleoyl-*sn*-glycero-3-phosphocholine (PC), 1,2-dioleoyl-sn-glycero-3-phospho-1’-*rac*-glycerol (PG), and 18: 1 cardiolipin (1’, 3’-bis[1,2-dioleoyl-*sn*-glycero-3-phospho]-*sn*-glycerol, sodium salt) (CL) were obtained from Avanti Polar Lipids (Alabaster, AL). Lipid mixtures at different molar ratios were dissolved in chloroform-methanol 2:1 (volume ratio), dried under nitrogen, and vacuum-lyophilized for 1 hour. After evaporation of the solvent, dried lipid films were hydrated by 20 mM Tris-HCl and 50 mM NaCl (TN buffer, pH 7.9) to a final lipid concentration of 5 mM. Small unilamellar vesicles (SUVs) were prepared by sonication of lipid suspension in cold water in a Branson M2800 ultrasonic bath sonicator for 20 minutes (Mendez, 2023), followed by passing them through 0.1 µm polycarbonate membranes (Avanti Polar Lipids) 13 times. SUVs can be stored at 4 °C for 7 days without freezing/thawing.

### Membrane flotation assay

The liposome flotation assays were performed according to Prochaska et al. with modification (Prochaska et al., 2018). Briefly, 1 mM SUVs were incubated with 5 µM purified PSB15 protein in 100 µl TN buffer for 1 hour at room temperature. In some experiments, 0.1 to 100 µM of GST-TrxA was added in SUVs-PSB15 mixtures. The liposome-bound and free PSB15 were separated by step sucrose gradient (2M, 0.75 M, 0 M) in 5 ml TN buffer centrifugation at 46,500 *g*, 22 °C for 30 min. The floated liposomes were collected and suspended with 5 ml TN buffer, then pelleted at 100,000 *g*, 22 °C for 30 min. The SUV-bound protein and input were analyzed by SDS-PAGE and immunoblotting.

For the fluorescent quantification of liposome-PSB15 binding, 5 µM GFP-PSB15 was incubated with 1 mM SUVs. For liposome-PSB15-DNA complex formation, 10 nM 5’-FITC-labeled PS oligonucleotide (IDT) was incubated with 1mM SUVs and 5 µM PSB15 mixtures and floated in a sucrose gradient as described above. To test TrxA effect on PSB15/DNA complex binding to SUVs, 0.1 to 100 µM of GST-thioredoxin was included in the mixtures prior to flotation. 50 µL of liposome fractions were collected and labeled with 10 µM amphiphilic styryl dye FM4-64 (ThermoFisher Scientific T13320) for 10 min at room temperature. The liposome was pelleted in TN buffer and resuspended with 200 µL TN buffer. The samples were loaded on a 96 well plate (BD), and the fluorescence intensity was read in a SpectraMax Gemini EM plate reader (Molecular Device) at 25 °C with settings as the following: GFP-PSB15 and FITC-labeled PS oligonucleotide (Excitation: 488 nm, Emission: 520 nm, cutoff: 515 nm), and FM4-64-labled SUVs (Excitation: 505 nm, Emission: 725 nm, cutoff: 630 nm). The fluoresce intensity of GFP-PSB15 and the FITC-PS oligonucleotide were normalized with FM4-64-labled SUVs. The apparent binding constant constants (K_d, app_) for the interaction of PSB15 with liposomes were calculated (Xu et al., 2012).

### Subcellular localization of PSB15

Cultures of Xcc-TcR transformed with pPSB15 plasmids were grown in LB medium up to an OD600 of 0.7. Periplasmic, cytoplasmic, inner, and outer membrane fractions were prepared as described (Hu et al., 1992; Chen et al., 1996). 20 μg of proteins from each fraction were analyzed by 12.5% SDS-PAGE. The OM and IM proteins were detected by immunoblotting using anti-OprF (ThermoScientific PA5-117553, 1:1000) and -MsbA (Abmart, X-P60752-C, 1:1000) antibodies, respectively (Doerrler et al., 2004; Woodruff and Hancock, 1989).

Xcc-TcR was infected with ϕLf-UK phage for 6 hours, and the IM and OM was analyzed on step sucrose gradient (25 to 61% [wt/wt]) centrifugation according to the procedures of Hu et al. (Hu et al., 1992). Sucrose densities of each fraction was determined from the refractive index (Cole-Parmer). The distribution of PSB15, MsbA, and OprF proteins were detected by immunoblotting the sucrose gradient fractions after chloroform/methanol (1:4) precipitation.

### Chromatin immunoprecipitation

pPSB15 mutant plasmids were transformed in Xcc-TcR. After grown at 28°C for 18 hours, bacteria were infected with ϕLf-UK-ϕPSB15 phage for 6 hours at 28°C. After IM was isolated as above, PSB15 protein and phage DNA were cross-linked with 1% formaldehyde, incubating for 15 min at 28°C. The cross-linking reaction was quenched by 0.125 M glycine and incubated for 5 min at 28°C. DNA was sheared with six cycles of 20 s pulses at 30% of maximum power with intermittent cooling on ice for 30 s using Branson’s M2800 ultrasonic bath sonicator. PSB15-DNA samples were incubated with 2 μg anti-PSB15 antibody and the mixture was rotated overnight at 4°C. The samples were mixed with 30 μl ChIP-grade protein A/G plus agarose beads (1: 1 slurry, ThermoFisher Scientific 26159) for 1 hour at 4°C. After several washes, the cross-linking was reversed by boiling and recovering from beads (Gade and Kalvakolanu, 2012). Phage DNA samples were amplified by PCR with primers (5’-GCCGCAGAGAAGATCCAAAAGAAC-3’ and 5’-GGAAATACCACGCACCACGGAAG-3’) and analyzed in 1% 0.5 × TBE agarose gel. The cytoplasmic pool of ϕLf-UK-ϕPSB15 phage RF DNA was PCR amplified as the control.

### Electrophoretic mobility shift assays (EMSA)

DNA oligonucleotides were synthesized by IDT. Binding between the 20 ng DNA oligonucleotides and 1 μg PSB15 proteins was carried out in a final volume of 15 μl in the DNA binding buffer (50 mM Tris-HCl pH 7.9, 50 mM NaCl, 1 mM EDTA, 1 mM BME, 10% (v/v) glycerol, 0.1 mg/ml BSA) for 30 min at 25°C. In the “supershift” experiment, 1 μg antibody was included in the binding reaction. 1.5 or 2 % agarose gel solutions (50 mL total volumes) were prepared with agarose powder (Bio-Rad) using 0.5× TBE buffer (45 mM Tris, 45 mM boric acid, 1 mM EDTA) and was then poured into a taped 7 ×10 cm tray. Samples were run in 0.5× TBE buffer at 150V for 10 or 15 mins. After electrophoresis, gels were stained in 50 mL SYBR Gold (ThermoFisher Scientific) in 0.5× TBE buffer (1:5000 dilution) for 40 minutes, followed by destaining in 200 mL 0.5× TBE buffer for 15 minutes with shaking. Gels were imaged in Molecular Imager ChemiDoc XR+ Imaging System and quantified with ImageLab software (Bio-Rad, Inc.). *K*_D,app_ values (apparent dissociation constant) were calculated as described (Ream et al., 2016).

### Total internal reflection fluorescence (TIRF) microscopy

pGII-transformed Xcc-TcR was grown until OD_600_=0.5 and infected by ϕLf-UK-GFP-PSB15 or PS(-)-GFP-PSB15 phage at a multiplicity of infection of 1 for 30 min. Bacterial IM was labeled by FM 4-64 (Fishov and Woldringh, 1999). 200 μl of bacteria was stained with 4 µM FM 4-64 in the medium for 20 min. 4 μl of bacteria was mixed with 80 μl melted agarose (1.5% BD AgarPlaque agarose in 1×phosphate-buffered saline), dropped on a Superfrost Plus precleaned glass slide (Fisher 1255015), and then immediately covered with a Gold Seal cover glass (24×60 mm, 0.13 mm thickness) (ThermoFisher Scientific). The samples were kept in the dark and at room temperature for 20 minutes before imaging. TIRF and epifluorescence images were acquired with an Olympus IX83 inverted microscope equipped with a cellTIRF-4Line module and an Olympus 1.49 numerical aperture 100× Uapo objective. The laser intensity settings on the main laser module were 30 mW for 488 nm (LuxX, Omicron, Germany). A fluorescence filter cube containing a polychroic beamsplitter (R405/488/561/647, Semrock) and a quad-band emission/blocking filter (FF01 446/523/600/695, Semrock) was used. Fluorescence images were captured at 1 Hz frame rates for 30 to 60 seconds using a Hammamatsu ImagEM X2 EM-CCD camera.

### Image analysis

The fluorescence emission intensity of each punctum was measured by the maximum intensity from the region of interest (ROI). The maximum intensity from each molecule was measured by first circling a ROI around the molecule. MetaMorph imaging software (Molecular Devices, Sunnyvale, CA) was used to automatically select the brightest pixel from each ROI and subtract the average background intensity from each image. The raw image data in each frame of the movie was analyzed with ImageJ using a GDSC SMLM plugin, which can determine the position of fluorescent molecules with sub-resolution precision (typically ∼<30 nm) (Etheridge et al., 2022). For dwell-time analysis, the start of each event in ROI was marked by an abrupt increase (> 10 A.U.) in the fluorescence signal, whereas the end of each event was marked by an abrupt decrease in the fluorescence signal (Asbury, 2016). Selecting the start and end of each event yielded the dwell time of each event, which was plotted in a histogram.

## ACKNOWLEDGMENTS

M.C.F. and P.J.F are scholarship recipients of the Olivia Constants Foundation. This work is inspired by Dr. Norton Zinder (1928-2012), David Pratt, Heinz Schaller, Robert Webster, Marjorie Russel, and Peter Model (1933-2017) and their lifetime achievement of filamentous phage assembly studies.

## DATA AVAILABILITY STATEMENT

The original contributions presented in the study are included in the article.

## Appendices

Figure S1 to S3, Table S1 to S3, and Movies S1 to S7 are included in Supplemental materials.

