## Supplemental figures, tables, movies and legend for "A packaging signal-binding protein regulates the assembly checkpoint of integrative filamentous phages"

**Appendices**

**Supplemental Figure Legend**

Figure S1. The flow chart of the suppressor screening of PS mutations. This two-step method utilizes the plaque morphology, followed by transducing particle (TP) production, to identify pseudorevertants that rescued phage assembly deficiency caused by T340A/C341G and the PS(-).

Figure S2. Samples are analyzed in 5-20% SDS-PAGE gel and immunoblotted with the affinity-purified anti-PSB15 antibody. Lane 1: recombinant PSB15 (20 ng, after enterokinase digestion), Lane 2: purified recombinant PSB15 (1 ng, after gel filtration), lane 3: CsCl purified ϕLf-UK virion (4 μg), lane 4: ϕLf-UK-infected Xcc-TcR (20 μg lysate), lane 5: Xcc-TcR infected with ϕLf-UK-ΔPSB15 phage (20 μg lysate), lane 6: pPSB15-transformed Xcc-TcR infected with ϕLf-UK-ΔPSB15 phage (20 μg lysate).

Figure S3. Gel filtration chromatography-purified recombinant PSB15 mutant proteins (4 μg) were analyzed in a 13.5 % SDS-PAGE gel and stained by Coomassie blue. Marker (M) is 16 kDa.

Figure S1

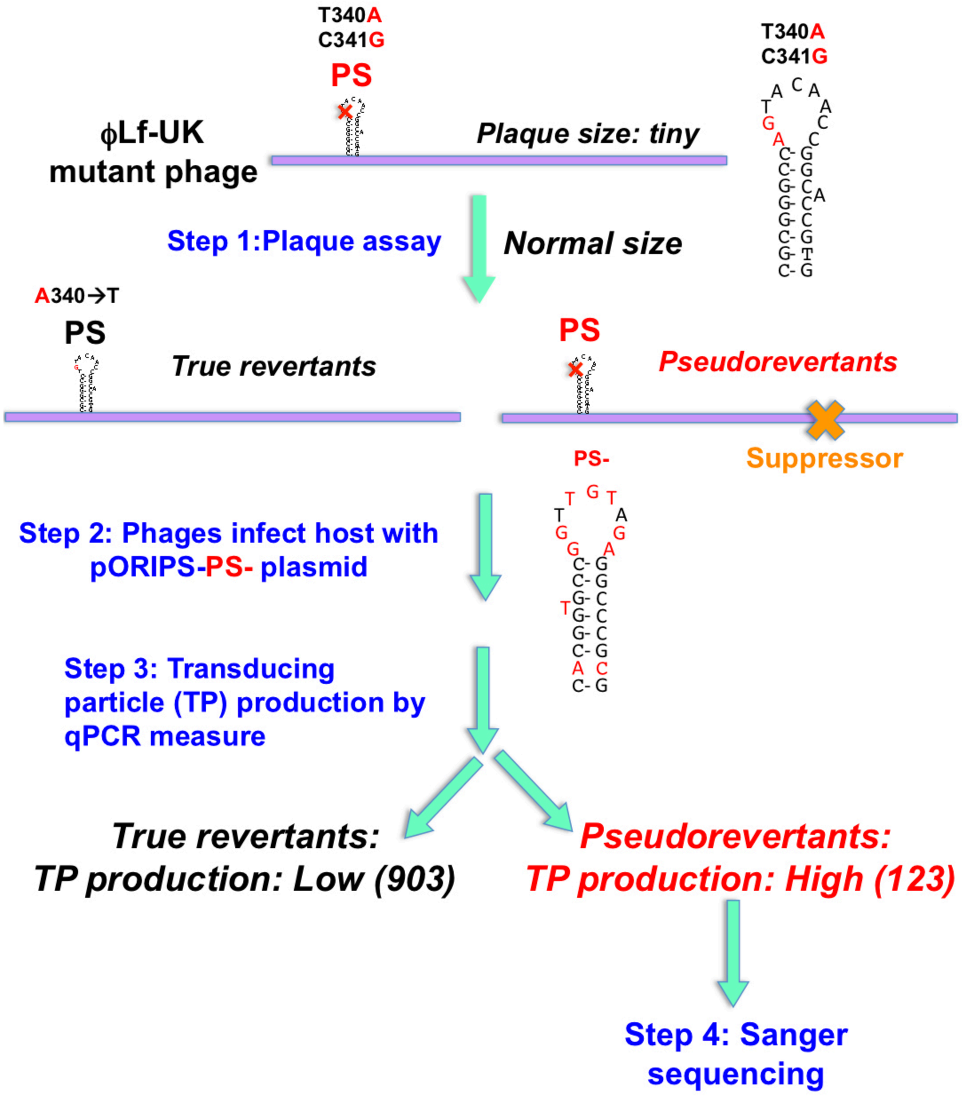

T340A
C341G
PS
φLf-UK
mutant phage
Plaque size: tiny
Step 1:Plaque assay
Normal size
A340→T
PS
True revertants
PS
Pseudorevertants
Suppressor
PS-
Step 2: Phages infect host with pORIPS-PS- plasmid
Step 3: Transducing particle (TP) production by qPCR measure
True revertants:
TP production: Low (903)
Pseudorevertants:
TP production: High (123)
Step 4: Sanger sequencing

Figure S2

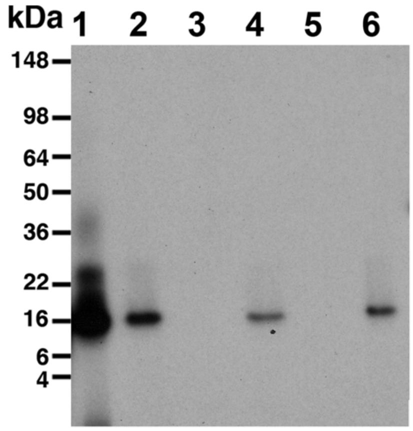

**Figure S3**

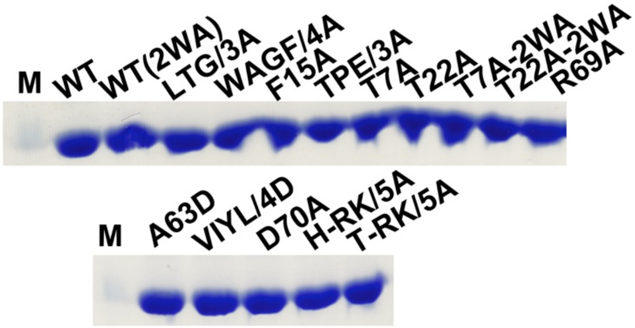
M
WT
WT(2WA)
LTG/3A
WAGF/4A
F15A
TPE/3A
T7A
T22A
T7A-2WA
T22A-2WA
R69A
M
A63D
VIYL/4D
D70A
H-RK/5A
T-RK/5A

**Supplemental Table Legend**

**Supplemental Table S1.** PS DNA-binding affinity of PSB15. The $K_{d, app}$ values unable to be detected (>50 μM) are labeled as "not available" ("N.A."). The PS sequences, mutations, and TP assembly efficiency (asterisk) were reported by Yeh et al. (24). The ones with low TP production yield are labeled in bold.

**Supplemental Table S2.** The PS DNA-binding affinity and the phage production yield of PSB15 mutations. The $K_{d, app}$ values unable to be detected (> 50 μM) are labeled as "not available" ("N.A."). ϕLf-UK-ΔPSB15 or PS(-)-ΔPSB15 RF DNAs was co-transformed into Xcc-TcR with pGII and pPSB15 as indicated. The phage DNA copy numbers released in the culture medium and the intracellular RF DNA replication were measured using qPCR as described (Yeh et al., 2023). The ones with low phage production yield are labeled in bold.

**Supplemental Table S3.** The binding affinity between FM4-64-labeled small unilamellar vesicles (SUVs, PE: PG: CL: PC=52%: 13%: 30%: 5%) and GFP-PSB15 as indicated. The $K_{d, app}$ values unable to be detected (>100 μM) are labeled as "not available" ("N.A.").

**Supplemental Table S1.**

| PS | DNA binding $K_{d,\,app}$ (nM) | TP assembly (%)* | PS Mutation |
|---|---|---|---|
| ϕLf-UK/ ϕLf | 10.2 ± 1.1 | 100 | |
| Xf109 | 12.7 ± 2.1 | 96-102 | |
| Cf1c/XacF1 | 10.9 ± 1.8 | 99-103 | |
| ϕLf2 | 11.3 ± 2.7 | 96-103 | |
| Cf2 | 11.5 ± 1.9 | 98-101 | |
| ϕXv2 | 12.2 ± 1.2 | 98-103 | |
| XaF13 | 13.4 ± 2.1 | 95-101 | |
| Xf409 | 12.2 ± 2.5 | 97-105 | |
| poly(A)$_{25}$ | N.A. | **0.5-1.2** | |
| poly(G)$_{25}$ | N.A. | **0.6-1.1** | |
| poly(T)$_{25}$ | N.A. | **0.7-1.1** | |
| poly(C)$_{25}$ | N.A. | **0.6-1.2** | |
| fLoop | N.A. | **1.1-1.5** | loop |
| T340A/C341G | N.A. | **0.9-1.4** | loop |
| C344G | N.A. | **0.9-1.3** | loop |
| PS(-) | N.A. | **0.9-1.5** | stem+loop |
| fSL | 12.8 ± 1.6 | 98-104 | stem+loop |
| fStem | N.A. | **1.2-1.6** | stem |
| C332A/G357T | N.A. | **1.1-1.5** | stem |
| G333C/T356G | N.A. | **0.8-1.3** | stem |
| G333A | N.A. | **1.2-1.5** | stem |
| T356C | 11.7 ± 2.1 | 98-101 | stem |
| C334A/G355T | N.A. | **1.1-1.4** | stem |
| G335A/C354T | N.A. | **0.9-1.3** | stem |
| G336A/C353T | N.A. | **1.0-1.5** | stem |
| G337T/C351A | 12.2 ± 2.3 | 96-103 | stem |
| Δloop | N.A. | **0.9-1.2** | loop |
| AT-loop | N.A. | **0.9-1.4** | loop |
| T-loop | N.A. | **1.0-1.6** | loop |
| A-loop | N.A. | **0.8-1.5** | loop |
| C-loop | N.A. | **1.1-1.4** | loop |
| T340A | N.A. | **1.1-1.3** | loop |
| C341G | 12.4 ± 2.8 | 95-101 | loop |
| T342G | 11.3± 2.6 | 98-104 | loop |
| A345G | 13.2 ± 2.3 | 97-102 | loop |
| A346G | 10.2 ± 2.5 | 96-101 | loop |
| C347G | 13.2 ± 2.5 | 98-102 | loop |
| C348G | 12.8 ± 1.8 | 97-103 | loop |

**Supplemental Table S2.**

| PS | PSB15 | DNA binding $K_{d,\,app}$ (nM) | Phage DNA copy number(/ml) |
|---|---|---|---|
| PS | WT | 10.2 ± 1.1 | $3.1\times10^{11} \pm 4.3\times10^{10}$ |
| PS | WT(2WA) | 11.3 ± 2.1 | $2.9\times10^{11} \pm 3.7\times10^{10}$ |
| PS | LTG/3A | N.A. | **$3.5\times10^{7} \pm 5.3\times10^{6}$** |
| PS | WAGF/4A | N.A. | **$2.7\times10^{7} \pm 6.6\times10^{6}$** |
| PS | F15A | N.A. | **$3.1\times10^{7} \pm 4.6\times10^{6}$** |
| PS | TPE/3A | N.A. | **$3.3\times10^{7} \pm 5.5\times10^{6}$** |
| PS | T7A | 10.8 ± 3.1 | $2.8\times10^{11} \pm 6.3\times10^{10}$ |
| PS | T22A | 12.7 ± 2.5 | $3.3\times10^{11} \pm 4.5\times10^{10}$ |
| PS | T7A(2WA) | 11.2 ± 2.8 | $3.0\times10^{11} \pm 5.2\times10^{10}$ |
| PS | T22A(2WA) | 10.5 ± 2.5 | $3.2\times10^{11} \pm 3.8\times10^{10}$ |
| PS | R69A | 12.6 ± 1.3 | $2.9\times10^{11} \pm 5.4\times10^{10}$ |
| PS | A63D | 11.3 ± 1.5 | **$3.1\times10^{7} \pm 3.6\times10^{6}$** |
| PS | VIYL/4D | 10.5 ± 1.1 | **$2.9\times10^{7} \pm 4.7\times10^{6}$** |
| PS | D70A | 12.8 ± 2.5 | **$3.2\times10^{7} \pm 5.1\times10^{6}$** |
| PS | H-RK/5A | 13.4 ± 1.8 | **$3.0\times10^{7} \pm 6.3\times10^{6}$** |
| PS | T-RK/5A | 12.3 ± 1.7 | **$2.8\times10^{7} \pm 3.6\times10^{6}$** |
| PS(-) | WT | 11.7 ± 2.2 | **$3.4\times10^{7} \pm 4.6\times10^{6}$** |
| PS(-) | WT(2WA) | 12.4 ± 2.4 | **$3.6\times10^{7} \pm 3.8\times10^{6}$** |
| PS(-) | T7A | 10.6 ± 2.9 | $2.9\times10^{11} \pm 5.3\times10^{10}$ |
| PS(-) | T22A | 12.1 ± 2.1 | $3.2\times10^{11} \pm 6.2\times10^{10}$ |
| PS(-) | T7A(2WA) | 13.9 ± 1.4 | $3.3\times10^{11} \pm 3.7\times10^{10}$ |
| PS(-) | T22A(2WA) | 10.1 ± 1.5 | $3.0\times10^{11} \pm 4.3\times10^{10}$ |

**Supplemental Table S3.**

| PSB15 | SUV binding $K_{d,\,app}$ (μM) |
|---|---|
| WT | 2.5 ± 0.4 |
| **WT(2WA)** | 2.4 ± 0.5 |
| **LTG/3A** | 2.7 ± 0.4 |
| **WAGF/4A** | 2.4 ± 0.6 |
| **F15A** | 2.3 ± 0.5 |
| **TPE/3A** | 2.6 ± 0.6 |
| **T7A** | 2.4 ± 0.4 |
| **T22A** | 2.5 ± 0.6 |
| **T7A(2WA)** | 2.5 ± 0.4 |
| **T22A(2WA)** | 2.3 ± 0.5 |
| **R69A** | 2.6 ± 0.3 |
| **A63D** | 2.5 ± 0.6 |
| **VIYL/4D** | 2.3 ± 0.4 |
| **D70A** | 2.7 ± 0.4 |
| **H-RK/5A** | N.A. |
| **T-RK/5A** | N.A. |

**Supplemental Movies Legend**

**Movies S1 to S7.** Representative movies of PSB15 dynamics shown in Figure 6A. pGII-transformed Xcc-TcR bacteria were infected with ϕLf-UK GFP-PSB15 (movies S1, WT/PS), ϕLf-UK GFP-PSB15-A63D (movies S2, A63D/PS), ϕLf-UK GFP-PSB15-F15A (movies S4, F15A/PS), PS(-)-GFP-PSB15 [movies S5, WT/PS(-)], ϕLf-UK GFP-PSB15-T7A (movies S6, T7A/PS), and PS(-)-GFP-PSB15-T7A [movies S7, T7A/PS(-)] phages at a multiplicity of infections of 1 for 30 min. Xcc-TcR TrxA G93D bacteria were transformed with pGII plamid and infected with ϕLf-UK GFP-PSB15 phage (movies S3, TrxA/G93D). All images were captured at 1 Hz frame rates for 30 to 60 seconds.
